# Spatial multi-omics of multiple myeloma uncovers niche-dependent pro-myeloma and immunosuppressive signaling in the bone marrow and extramedullary lesions

**DOI:** 10.64898/2026.03.23.713195

**Authors:** Denis J Ohlstrom, Marina E Michaud, Mojtaba Bakhtiari, Junia Vieira Dos Santos, William C Pilcher, Alden Staub, Sarthak Satpathy, Katherine Ferguson, Sowmitri Mantrala, Seunghee Kim-Schulze, Zhihong Chen, Sagar Lonial, Melissa L Kemp, Daniel Sherbenou, Alessandro Lagana, David L Jaye, Ajay K Nooka, Samir Parekh, Manoj K Bhasin

**Affiliations:** Coulter Department of Biomedical Engineering, Georgia Institute of Technology, Atlanta, GA, USA; Department of Pediatrics, Emory School of Medicine, Atlanta, GA, USA; Tisch Cancer Center, Department of Hematology and Oncology, Precision Immunology Institute, Genetics and Genomic Sciences, Icahn School of Medicine at Mount Sinai, New York, NY, USA; Department of Biomedical Informatics, Emory School of Medicine, Atlanta, GA, USA; Tisch Cancer Institute, Department of Immunology and Immunotherapy, Human Immune Monitoring Center (HIMC), Icahn School of Medicine at Mount Sinai, New York, NY, USA; Aflac Cancer and Blood Disorders Center, Children’s Healthcare of Atlanta, Atlanta, GA, USA; University of Colorado Denver-Anschutz Medical Campus, Aurora, CO; Department of Pathology and Laboratory Medicine, Emory University, Atlanta, GA 30322, USA; Department of Hematology Oncology, Emory School of Medicine, Atlanta, GA, USA; Division of Nephrology, The Ohio State University Wexner Medical Center, Columbus, Ohio, USA

**Keywords:** Multiple Myeloma, Immune Microenvironment, Risk Stratification, Single-Cell, Transcriptome, Exhaustion

## Abstract

Multiple myeloma (MM) is a plasma cell malignancy shaped by dynamic interactions between MM cells and non-malignant cells in the immune microenvironment. To spatially profile the influence of cellular context on MM and immune cell expression, we developed a multimodal framework integrating 10x Genomics Visium HD, 10x Genomics Xenium, and clinically annotated single-cell RNA (scRNA-seq) sequencing datasets. Visium HD enabled unbiased, whole transcriptome, spatial discovery at 16 µm resolution, Xenium provided orthogonal validation at single-cell resolution, and scRNA-seq extended findings by mapping spatial labels and leveraging the greater sequencing depth. We developed a custom framework for cell type annotation within Visium HD spatial bins. Our approach enabled identification of plasma cell-dense niches enriched for non-canonical Wnt signaling, associated with gene expression supporting cell adhesion mediated drug resistance, inferior progression-free survival, and extramedullary lesions. Immune cells within these neighborhoods exhibited suppressed transcriptional states, including increased inhibitory receptor expression such as *LAG3*. Utilizing the niche-driven transcriptional states in MM and immune cells, we were able to develop a 15-gene signature independently predictive of progression free survival (HR = 2.00, p < 0.0001). Collectively, this study demonstrates the potential of integrated spatial and single-cell transcriptomics to define niche-specific programs supporting MM progression.

## Introduction

Multiple myeloma (MM), the second most common hematoogic malignancy, is a plasma cell cancer characterized by near-universal disease relapse(1). MM incidence has risen to an estimated 36,000 new diagnoses annually in the United States, with approximately 13,000 deaths per year(1). Although advances in MM-targeted therapies, autologous stem cell transplantation (ASCT), and immunotherapy have improved 5-year survival rates(2), most patients ultimately experience disease progression(3,4). Treatment response and disease evolution are influenced by multiple factors, including cytogenetic abnormalities, intra-tumoral heterogeneity, and the cellular and molecular composition of the bone marrow microenvironment (BMME)(5,6). High-dimensional profiling approaches, particularly single-cell RNA sequencing (scRNA-seq), have provided important insights into the transcriptional programs of malignant plasma cells and non-malignant BMME compartments associated with therapeutic resistance and early progression(7–10). However, these methods require tissue dissociation, disrupting native spatial architecture and obscuring critical cellular interactions within microenvironmental niches that influence tumor behavior.

Recent technical advances in spatial transcriptomic and proteomic profiling have begun to address this limitation. Approaches such as laser capture microdissection coupled with sequencing(11,12) or fluorescence-based assays such as CODEX (CO-Detection by indEXing)(13,14) have revealed the structured spatial organization of malignant and immune populations within the BMME at cellular resolution. The introduction of sequencing-based spatial transcriptomic platforms such as 10x Genomics Visium further enabled genome-wide expression profiling at ∼55 µm resolution, revealing regional transcriptional heterogeneity of malignant and immune cells within MM lesions(12,15). Despite these advances, current technologies require a tradeoff between spatial resolution and molecular breadth: higher-resolution platforms typically rely on targeted gene panels, whereas unbiased transcriptome-wide approaches operate at lower spatial resolution. A recent landmark study by Yip et al.(16) significantly advanced the field by applying the fluorescence-based 10x Genomics Xenium spatial platform to MM bone marrow (BM), enabling high-resolution mapping of immune and malignant cell phenotypes and delineation of lymphoid, myeloid, and stromal compartments *in situ*. While these innovations have substantially improved our ability to define cellular composition and spatial architecture, they remain limited in their capacity to perform unbiased, spatially resolved differential expression analyses across the full transcriptome with cellular resolution. Thus, approaches that integrate high spatial resolution with comprehensive transcriptomic profiling are still needed.

To achieve this goal, we developed an integrated multimodal framework combining sequencing-based spatial transcriptomics, fluorescence-based spatial transcriptomics, and scRNA-seq. For unbiased, whole-transcriptome discovery, we profiled 21 MM BM biopsies using the 10x Genomics Visium HD (V-HD) spatial transcriptomics platform, enabling high-resolution spatial profiling of the BMME at 16 µm resolution. To complement the high transcriptomic complexity captured by V-HD with true single-cell resolution, we orthogonally validated our findings using the spatial transcriptomics data generated by Yip et al. using the 10x Genomics Xenium platform(16). Finally, we mapped the spatial gene expression signatures onto myeloma and immune cell populations using the clinically annotated scRNA-seq dataset published by Pilcher et al.(8), leveraging the large cohort size and greater transcriptomic depth to enable more comprehensive downstream characterization. Together, this multimodal strategy integrates spatial context with single-cell transcriptomic depth, providing a robust and high-resolution framework to dissect the cellular architecture and molecular programs of the MM bone marrow microenvironment.

## Results

### Construction of a spatial atlas of multiple myeloma

To interrogate the spatial interactions between MM and immune cells in the BMME, we combined complementary transcriptomic platforms to generate a multimodal spatial atlas. In this framework, we utilized V-HD for whole-transcriptome, discovery-driven profiling, Xenium for orthogonal validation at single-cell spatial resolution with reduced gene coverage, and scRNA-seq for high-depth transcriptomic capture to extend our spatial findings (**Fig. 1A**). For V-HD, twenty-one MM bone marrow core biopsies were profiled (**Supplemental Fig. 1**). Samples were selected to cover a broad range of recurrent cytogenetic aberrations including eight with odd numbered trisomies, ten with t(4;14), five with t(11;14), four with gain(1q), four with del(17p), and ten with del(13q) (**Fig. 1B**). Sixteen of the biopsies were collected at disease diagnosis and five nonlongitudinal biopsies were collected at disease relapse. Biopsies ranged in histologically estimated plasma cell percentage from 5 to 90%.

**Figure 1.**
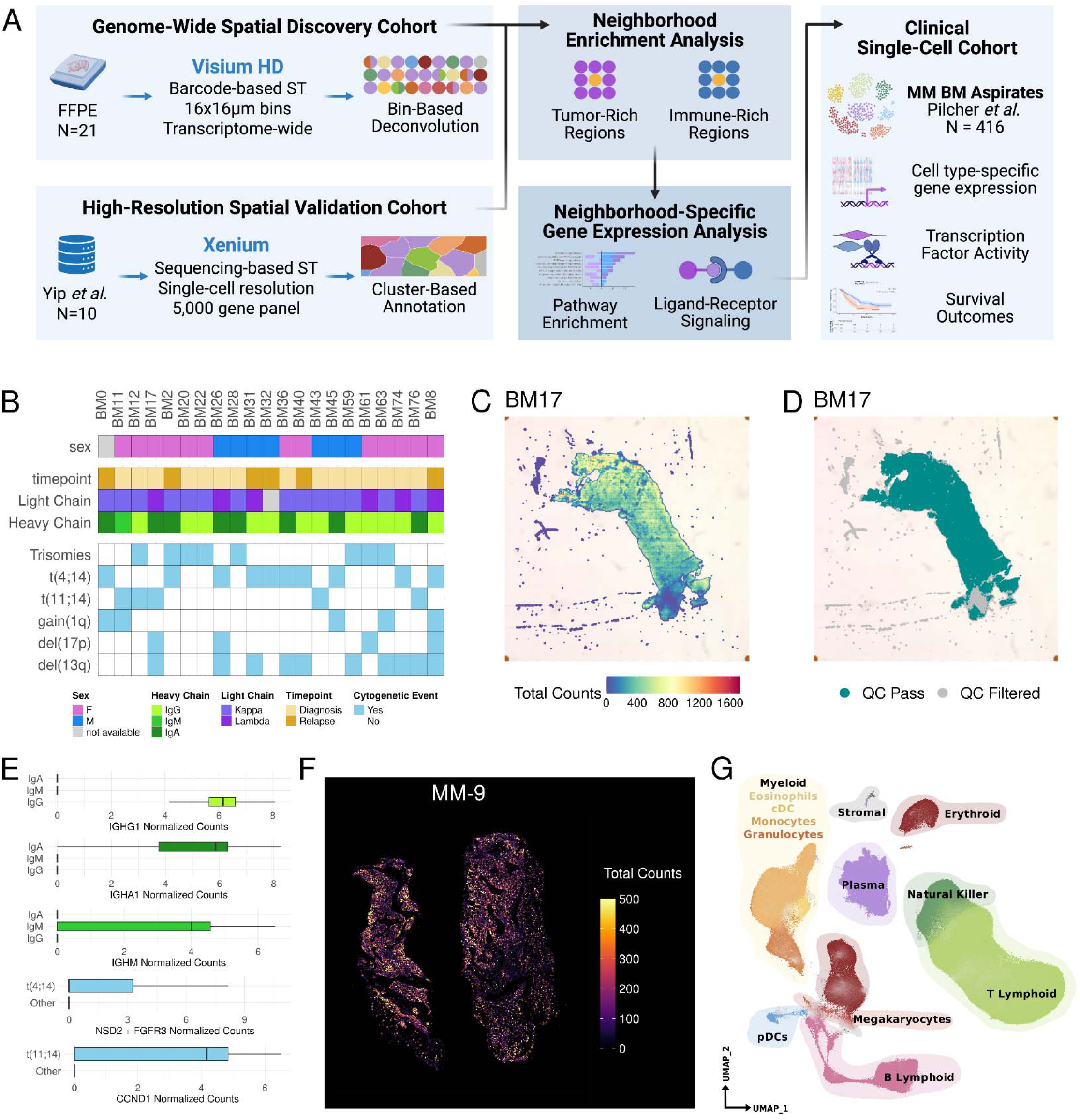
Construction of a spatial atlas of multiple myeloma. **(A)** Schematic overview of the analytical approach. Visium-HD spatial transcriptomics was used for whole-genome discovery-driven analyses, Xenium spatial transcriptomics was used for single-cell resolved orthogonal validation, and single-cell RNA sequencing was used for high depth transcriptomic analyses. **(B)** Metadata heatmap showing the clinical and cytogenetic features of the samples profiles by Visium-HD. **(C)** Representative spatial dot plot displaying the total gene detection per voxel for sample BM17. See Supplemental Figure 2 for quality control metrics across samples. **(D)** Representative spatial dot plot for voxels passing quality control (teal) or filtered out from downstream analyses (gray). Quality control cutpoints were set as total counts ≥ 100 and total mtRNA ≤ 20%. **(E)** Box plots displaying the gene expression of immunoglobulin heavy chains and genes involved in recurrent translocation split by each patients’ involved immunoglobulin and presence/lack of recurrent translocation. **(F)** Representative spatial dot plot displaying the total gene detection per voxel for sample MM-9 from Yip et al., the Xenium dataset used for orthogonal validation. See Supplemental Figure 3 for additional quality control metrics. **(G)** Uniform manifold approximation and projection plot displaying cell types in the single-cell RNA sequencing dataset published by Pilcher et al. used for high depth analytical extension of the spatial findings. See Supplemental Figure 4 for cell type annotations.

A total of 252,249 16 x 16 µm voxels passed quality control filters (≥100 unique molecular identifiers, <20% mtRNA) with an average of 289.6 counts and 194.2 unique genes per voxel (**Fig. 1C-D, Supplemental Fig 2A**). Transcriptome coverage was reasonably consistent across samples, with an average of 16,164 unique genes (range of 12,235 to 18,041) detected per biopsy (**Supplemental Fig. 2B**). Total voxel detection varied between samples from 1,545 to 32,665 (**Supplemental Fig 2C**). Transcript detection of the involved immunoglobulin heavy chain (e.g., *IGHG1* in patients with IgG MM) and genes amplified by recurrent translocations (e.g., *CCND1* in t(11;14)) were robustly detected, indicating successful capture of MM gene profiles (**Fig. 1E**).

While 16 x 16 µm voxels in V-HD provide whole-transcriptome quantification at near single-cell resolution, each voxel can encompass transcripts from multiple adjacent cells due to the fixed grid and voxel size. To address this limitation, we incorporated two complementary, publicly available datasets into our atlas. First, we analyzed ten MM bone marrow biopsies recently profiled by Yip *et al*. using Xenium (**Fig. 1F, Supplemental Fig. 3A-F**)(16). Although the Xenium assay targets a curated panel of ∼5,000 genes rather than the full transcriptome, it provides single-cell resolution. We therefore used the Xenium data to provide orthogonal validation of Visium-derived findings for genes shared between the two platforms. Second, we included single-cell RNA sequencing (scRNA-seq) profiles from 416 biopsies collected at diagnosis or relapse from 336 patients in the MM Immune Atlas, a dataset generated by multiple groups, including ours, in collaboration with the Multiple Myeloma Research Foundation (MMRF)(8) (**Fig. 1G, Supplemental Fig. 4A-C**). In contrast to spatial assays, scRNA-seq typically captures thousands of genes per cell, enabling deeper characterization of transcriptional programs and signaling pathways. Furthermore, the paired outcome data in this large, clinically annotated single-cell dataset facilitated robust validation of the prognostic associations of genes of interest. Together, the integration of V-HD, Xenium, and scRNA-seq datasets provided complementary transcriptome breadth, sequencing depth, and cellular resolution, strengthening the robustness of our analyses while enabling characterization of MM and immune transcriptional programs.

### Voxel deconvolution identifies expected cell types in the bone marrow microenvironment

To identify the cell type(s) contributing to the transcriptomic signal of each 16 x 16 µm voxel, we developed a spatial deconvolution framework tailored to V-HD data. Briefly, V-HD profiles were deconvolved against a reference scRNA-seq dataset (**Supplemental Fig. 5A-C**) using Robust Cell Type Decomposition (RCTD) algorithm(17), followed by assignment of contributing cell type(s) using an empirical Bayesian shrinkage approach (see Methods, **Fig. 2A**). This strategy enabled assignment of primary contributing cell type(s) to 198,717 voxels, while 28,627 voxels failed to meet threshold criteria for any cell type and were classified as “Mixed” (**Fig. 2B-C; Supplemental Fig. 6A**). The proportion of voxels annotated as plasma cell-containing correlated strongly with histologically estimated bone marrow plasma cell burden, supporting the plausibility of the annotations (**Supplemental Fig 6B**). To further assess the quality of annotations, we performed one-versus-all differential expression for each cell type. Annotated voxels consistently depicted significantly higher expression of canonical lineage markers, including *SDC1* in plasma cells (Log_2_ Fold Change [L_2_FC] = 2.48, multiple comparisons adjusted p value [adj.p] = 2.14×10^-87^), *CD3D* in T cells (L_2_FC = 4.94, adj.p = 4.90×10^-49^), and *MPO* in granulocytes (L_2_FC = 3.30, adj.p = 1.0×10^-250^, **Supplemental Fig. 6C**). The enrichment of canonical lineage genes demonstrates that our deconvolution framework accurately resolves the cell-type identities contributing to each voxel’s transcriptome, providing robust annotations for downstream spatial analyses.

**Figure 2.**
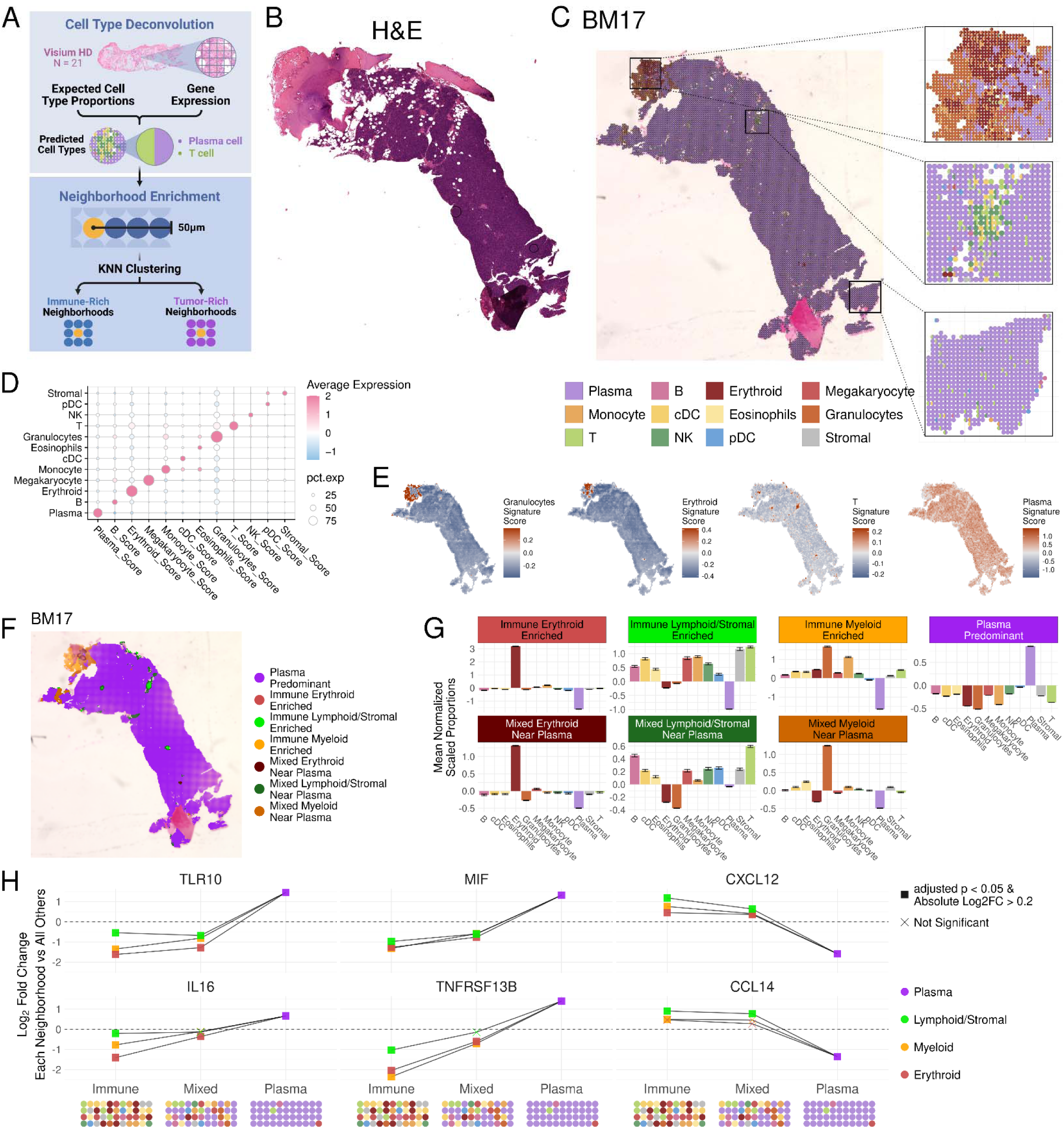
Voxel deconvolution identifies expected cell types in the bone marrow microenvironment. **(A)** Schematic overview of the voxel deconvolution approach. Voxels were deconvolved using a reference single-cell RNA sequencing dataset and voxels binarized as containing or not containing a given cell type using an empirical Bayesian shrinkage approach. Then voxels were grouped into neighborhoods based on the composition of proximal cells. **(B)** Hematoxylin and eosin staining of the bone marrow biopsy for BM17, see Supplemental Figure 1 for other stained sections. **(C)** Representative spatial scatterpie plot showing the annotation of voxels in BM17. Voxels are colored by cell type, when multiple cell types are present in a given voxel the scatterpie plot is equally split and colored by the cell types present. **(D)** Dot plot displaying the lineage marker gene scores for the cell types detected in the Visium HD dataset. Lineage marker gene scores were calculated as the scaled average normalized expression across 8-12 lineage marker genes per cell type (see Methods for gene lists). **(E)** Representative spatial hexbin plots displaying the density of lineage marker gene scores for BM17. **(F)** Representative spatial dot plot displaying the neighborhood annotations for BM17 after unsupervised K-nearest neighbor neighborhood analysis. See Supplemental Figure 8 for plots supporting the neighborhood process. **(G)** Bar plots depicting the relative abundance of cell types in the seven annotated neighborhoods. **(H)** Dot plot displaying the relative expression of selected ligand and receptor genes across the seven annotated neighborhoods. One-versus-all differential was performed for each neighborhood, the Y axis displays the resulting Log_2_ fold change and the dot shape indicates significance. Differential expression was tested using a hurdle model as implemented by the MAST method in Seurat’s FindMakers function with a significance cutoff of multiple comparisons adjusted p value < 0.05. See Supplemental Figure 11 for additional plots displaying the differential expression of ligands and receptors across neighborhoods.

Broad detection of abundantly expressed genes, notably the immunoglobulin heavy and light chain genes, is a known limitation of sequencing-based spatial transcriptomics of the BM, even in non-MM contexts where plasma cells are expected to be relatively sparse(15,18). Consistent with these prior projects, immunoglobulin genes were detected nearly ubiquitously in our data, even in biopsies where plasma cells accounted for less than 25% of all cells based on histology (**Supplemental Fig. 7A**). Our deconvolution framework mitigated these ubiquitous signals by leveraging the diversity of genome-wide transcriptional profiles, thereby improving cell-type specificity (**Supplemental Fig 7B**). As a result, annotated voxels exhibited selective enrichment of canonical lineage gene signatures, indicating that voxel assignment was driven by lineage-specific programs rather than individual broadly detected genes (**Fig. 2D-E, Supplemental Fig. 7C-E**). Together, these results indicate that the deconvolution framework robustly identifies the dominant cellular contributors to each voxel’s transcriptomic profile despite known limitations of the V-HD platform, enabling reliable downstream spatial analyses.

### Neighborhood annotation identifies cellular sub-niches within the bone marrow microenvironment

To characterize how local cellular composition influences gene expression, we next defined spatial neighborhoods by clustering voxels based on the composition of surrounding cell types. Voxels were grouped using unsupervised K-nearest neighbors (KNN) clustering on the proportion of cell types within a 50 µm radius of each voxel (**Supplemental Fig. 8A-B**). Resulting clusters were annotated as cellular neighborhoods based on over-represented cell types (**Supplemental Fig. 8C-D**). Seven compositional neighborhoods were identified in V-HD data: one plasma cell-predominant neighborhood, three neighborhoods enriched for non-malignant cells (lymphoid/stromal-predominant, erythroid-predominant, and myeloid-predominant), and three neighborhoods comprising mixtures of malignant and non-malignant cells (**Fig. 2F-G**). To evaluate cross-platform concordance, individual cells in the Xenium dataset were harmoniously annotated using the same cell-type labels as the V-HD dataset (**Supplemental Fig. 9A-D**), followed by the same KNN-based compositional analyses to elucidate spatial neighborhoods. Notably, the same neighborhoods were also observed in Xenium data (**Supplemental Fig. 10A-E**), suggesting that these cellular arrangements represent recurrent spatial sub-niches within the MM BMME.

Next, to characterize the signaling landscape within each spatial niche, we compared the expression of ligand and receptor genes between neighborhoods. The plasma cell-predominant neighborhood was enriched for ligands and receptors known to support MM survival, including *TNFRSF13B*(19) (L_2_FC = 1.40, adj.p = 7.39×10^-304^), *TLR10*(20) (L_2_FC = 1.45, adj.p = 1.14×10^-49^) and *IL16*(21,22) (L_2_FC = 0.66, adj.p = 5.23×10^-112^, **Fig. 2H**, **Supplemental Fig. 11A**). Plasma cell-predominant neighborhoods were also enriched for microenvironment-modulating molecules including *MIF* (L_2_FC = 1.32, adj.p = 2.90×10^-150^), which has been implicated in promoting T cell exhaustion in MM(23), as well as the pro-angiogenic genes *VEGFB* (L_2_FC = 0.33, adj.p = 2.70×10^-110^) and *CD320*(24) (L_2_FC = 0.85, adj.p = 1.79×10^-56^). In contrast, the immune-enriched and mixed neighborhoods exhibited broadly shared signaling features, with enrichment of factors such as *CXCL12* and *CCL14* observed across all six groups (**Supplemental Fig. 11B-G**). These findings suggest that the transcriptional profiles associated with each spatial neighborhood delineate the plasma cell-predominant niche from the remaining neighborhoods, rather than differentiating among the immune and mixed niches themselves.

To systematically evaluate the influence of spatial context on cell-intrinsic transcriptional programs, we performed principal component analysis (PCA) for each cell type across neighborhoods (e.g., T cell-containing voxels across all seven neighborhoods). Consistent with the ligand/receptor analysis, cells residing in immune and mixed neighborhoods shared similar transcriptomes, whereas cells in plasma cell-predominant regions typically exhibited distinct transcriptional profiles (**Supplemental Fig. 12A-L**). Independent neighborhood PCA in the Xenium dataset identified the same pattern, with cells in plasma-rich neighborhoods typically appearing distinct from the other six neighborhoods (**Supplemental Fig. 13A-K**). Collectively, these analyses indicate that the dominant axis of spatial transcriptional variation distinguishes the plasma cell-predominant niche from all other neighborhoods, leading us to further investigate this niche in greater detail.

### Plasma cells in plasma-rich neighborhoods are enriched for genes associated with chemoresistance, quiescence, and altered paracrine signaling

To define the transcriptional programs associated with plasma cell-dense regions of the bone marrow, we compared the gene expression in plasma cells from plasma cell-predominant neighborhoods versus plasma cells in non-plasma-dominant regions (**Fig. 3A**). Utilizing our V-HD data, we found 1,687 genes were differentially expressed between neighborhoods, with 249 differentially expressed in the same direction in the Xenium dataset, 1,229 not detected in Xenium, and 209 genes were either not differentially expressed or were differentially expressed in the opposite direction in Xenium (**Fig 3B**). Genes demonstrating concordant spatial differential expression across both platforms, together with V-HD-specific genes not detected in Xenium, were retained for downstream analyses. This approach prioritized reproducible niche-associated transcriptional programs while preserving the broader transcriptomic resolution uniquely afforded by V-HD.

**Figure 3.**
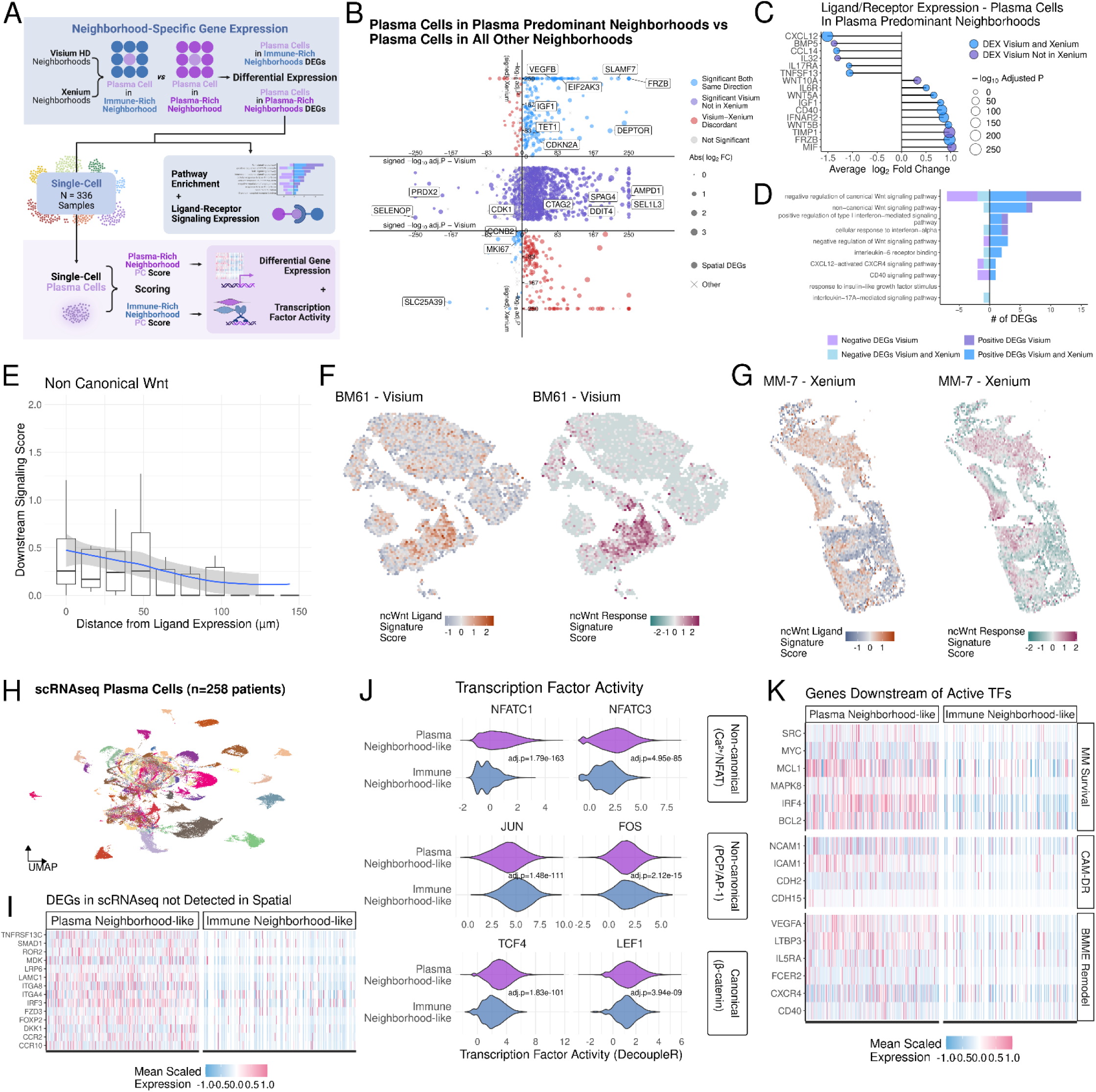
Plasma cells in plasma cell-predominant neighborhoods are enriched for non-canonical Wnt signaling. **(A)** Schematic overview of the spatial analysis framework. Plasma cells were compared between plasma cell-predominant neighborhoods and immune-enriched neighborhoods to find spatially differentially expressed genes. Spatially differentially expressed genes were then used to assign spatial labels to single-cell RNA sequenced plasma cells for higher depth transcriptomic analyses. **(B)** Dot plot displaying the independent differential expression of plasma cells in plasma cell-predominant neighborhoods versus plasma cells in immune enriched neighborhoods in both Visium HD and Xenium data. Axes display the -log_10_ adjusted p value signed by the log_2_ fold change (i.e. a gene with a negative log_2_ fold change is displayed on the negative portion of the graph). Genes displaying concordant differential expression (i.e. adjusted p value < 0.05 and log_2_ fold change with the same sign) are displayed in blue, while discordant expression is displayed in red. Given that Visium is a whole-transcriptome assay, many genes in the Visium data were not detected in Xenium data. Genes that were spatially differentially expressed in Visium but not detected in Xenium are in the center of the graph in purple. Differential expression was tested using a hurdle model as implemented by the MAST method in Seurat’s FindMakers function with a significance cutoff of multiple comparisons adjusted p value < 0.05. **(C)** Lollipop plot displaying spatially differentially expressed ligand and receptor genes. The X axis shows log_2_ fold change, lollipop size shows -log_10_ adjusted p value, and color indicates whether the ligand was differentially expressed in Visium and Xenium or Visium and not detected in Xenium. **(D)** Bar plot showing the number of differentially expressed genes present in downstream Gene Ontology Biological Pathways (GO:BP) of the differentially expressed ligands. Bars are colored by the assays in which differential expression was detected, negative values indicate genes with negative log_2_ fold change values. **(E)** Box plot displaying the relationship between downstream non-canonical Wnt signaling scores for plasma cells based on their distance from the nearest non-canonical Wnt ligand-expressing voxel. Downstream signaling score is calculated from genes present in GO:2000052 “non-canonical Wnt signaling pathway” and GO:00900090 “negative regulation of canonical Wnt signaling pathway”. **(F-G)** Representative spatial hexbin plots showing the localization of non-canonical Wnt ligand expression (left panels) and downstream signaling scores (right panels) for Visium (F) and Xenium data (G). Color scale represents relative density of ligand expression (blue-white-orange) and relative density of downstream signaling expression (green-white-pink). **(H)** Uniform manifold approximation and projection plot displaying the single-cell RNA sequenced plasma cells colored by patient. See Supplemental Figure 14 for mapping of spatial labels onto single-cell RNA data. **(I)** Heatmap displaying the mean normalized scaled expression of ligand and receptor genes that were differentially expressed between plasma-neighborhood-like plasma cells versus immune-neighborhood-like plasma cells and not well detected in spatial assays. **(J)** Violin plots of transcription activity inferred using DecoupleR. Transcription factors are grouped by row based on the Wnt pathway they are downstream of. **(K)** Heatmap displaying the mean normalized scaled expression of genes that are downstream of the differentially active transcription factors shown in (J).

Plasma cells in plasma cell-predominant neighborhoods exhibited transcriptional programs associated with disease persistence and microenvironmental adaptation. These cells differentially expressed genes associated with survival, chemoresistance, and unfolded protein stress durability, including *DEPTOR*(25) (L_2_FC = 1.66, adj.p = 1.49×10^-206^), *DDIT4*(26) (L_2_FC = 0.80, adj.p = x10^-148^), and *EIF2AK3*(27) (L_2_FC = 0.85, adj.p = 6.01×10^-92^), respectively. In parallel, these cells demonstrated increased expression of secreted ligands known to promote MM survival and BMME remodeling, including the growth factor, *IGF1*(28), (L_2_FC = 0.80, adj.p = 3.85×10^-19^), the canonical Wnt antagonist, *FRZB*(29) (L_2_FC = 0.98, adj.p = 1.32×10^-250^), and the pro-angiogenic factor, *VEGFB* (L_2_FC = 0.22, adj.p = 6.61×10^-69^). Collectively, these findings suggest that plasma cells in plasma cell-predominant neighborhoods adopt a survival-optimized state while actively reshaping the local signaling milieu. Given the prominence of secreted ligands among the top spatially enriched genes, we next interrogated the signaling landscape within plasma cell-predominant regions of the BMME.

### Plasma cells in plasma cell-predominant neighborhoods utilize auto/paracrine non-canonical Wnt signaling

To evaluate how autocrine and paracrine signaling programs distinguish plasma cells in plasma cell-predominant niches, we evaluated the differential expression of genes encoding secreted ligands and surface receptors relative to other niches. Plasma cells in plasma cell-predominant regions upregulated several notable mediators of Wnt signaling, including the canonical Wnt antagonist *FRZB* (L_2_FC = 0.98, adj.p = 1.32×10^-250^), the non-canonical Wnt agonists *WNT5B* (L_2_FC = 0.95, adj.p = 1.78×10^-27^) and *WNT5A* (L_2_FC = 0.66, adj.p = 3.81×10^-24^), and to a lesser extent the canonical Wnt agonist, *WNT10A* (L_2_FC = 0.98, adj.p = 1.32×10^-250^, **Fig. 3C**). Additionally, signaling molecules with established roles in shaping the BMME, namely *MIF* (L_2_FC = 1.02, adj.p = 1.38×10^-106^), *IFNAR2* (L_2_FC = 0.86, adj.p = 1.14×10^-152^), *IL32* (L_2_FC = −1.31, adj.p = 1.01×10^-10^), and *CXCL12* (L_2_FC = −1.51, adj.p = 1.00×10^-250^) were also differentially expressed. Notably, many of these ligands have been implicated in the promotion of cell adhesion-mediated drug resistance (CAM-DR) in MM, including *WNT5B*, *WNT5A*(29), and *MIF*(30), drawing a plausible connection between the enriched expression of genes for secreted ligands and the formation of plasma-dense niches in the BMME.

While many of the identified ligands in plasma cell-predominant niches have been reported to influence MM phenotypes, many also have known impacts on non-malignant cells in the BMME(23,31–34). To determine which ligands were most likely influencing plasma cell expression through autocrine signaling, we evaluated their potential downstream effects by examining the number of differentially expressed genes that correspond to ligand-stimulated Gene Ontology Biological Pathways (GO:BP)(35). The largest number of differentially expression genes was observed in the “negative regulation of canonical Wnt signaling” and “non-canonical Wnt signaling” pathways, consistent with elevated expression of *FRZB*, *WNT5B*, and *WNT5A* (**Fig. 3D**). Other pathways such as “CD40 signaling pathway” and “response to insulin-like growth factor stimulus” had comparatively fewer genes that were differentially expressed in plasma cell-predominant niches, potentially indicating that the ligands corresponding to these pathways are interacting with receptors on non-malignant cells in the BMME or representing a technical byproduct of the sparsity of current spatial assays. Given that the non-canonical Wnt signaling pathway exhibited coherent expression of both ligands and downstream signaling genes within the plasma cell-predominant neighborhoods, we next evaluated whether this paracrine signaling interaction could be validated by the spatial proximity of the ligand and its downstream gene expression.

To assess the spatial plausibility of the observed non-canonical Wnt signaling, we compared the expression of downstream non-canonical Wnt pathway genes with the distance to the nearest ligand-expressing voxel (i.e., canonical Wnt antagonist and non-canonical Wnt agonist). Voxels within 16 µm of ligand expression exhibited the highest downstream signaling scores, whereas voxels ≥100 µm away showed near-zero expression of downstream genes (**Fig. 3E**), demonstrating clear spatial dependency. Applying the same analysis to the Xenium dataset independently validated this spatial association between ligand expression and downstream pathway activation (**Fig. 3F-G**). Based on these findings, the spatial coupling between ligand proximity and downstream transcriptional activation is consistent with non-canonical Wnt signaling enrichment in plasma cell-predominant neighborhoods of the BMME.

### Wnt signaling supports expression of genes associated with MM survival, CAM-DR, and BMME remodeling

Given that both canonical and non-canonical Wnt agonists and antagonists were spatially differentially expressed, we next sought to determine which downstream transcriptional programs were preferentially activated in plasma cells from plasma cell-predominant neighborhoods. To enable higher-resolution genomic interrogation across a larger cohort, we leveraged our scRNA-seq dataset encompassing 96,880 plasma cells from 258 patients (**Fig. 3H**). Using the spatially differentially expressed genes as a neighborhood-defining signature, we identified plasma cells in the scRNA-seq dataset with transcriptomic profiles resembling those observed in plasma cell-predominant neighborhoods (**Supplemental Fig. 14A**). As expected, the plasma cells in scRNA-seq with plasma-neighborhood-like transcriptomes recapitulated key spatial findings, including increased *WNT5B* (L_2_FC = 1.21, adj.p = 5.3×10^-11^), *FRZB* (L_2_FC = 1.13, adj.p = 9.91×10^-45^), and *IFNAR2* (L_2_FC = 0.70, adj.p = 2.3×10^-88^) expression (**Supplemental Fig. 14B-C**). Furthermore, several genes that were not well detected in spatial assays were differentially expressed by plasma cell-predominant-like cells including the non-canonical Wnt receptor, *FZD3* (L_2_FC = 1.09, adj.p = 9.30×10^-40^), the non-canonical Wnt receptor, *ROR2* (L_2_FC = 1.45, adj.p = 2.0×10^-15^), and the canonical Wnt co-receptor, *LRP6* (L_2_FC = 1.72, adj.p = 1.16×10^-19^, **Fig. 3I**), providing additional evidence that altered Wnt signaling is central to the identity of plasma cells in plasma cell-predominant neighborhoods.

To more precisely define which Wnt signaling branches were functionally active in plasma-neighborhood-like cells, we next interrogated transcription factor activity to distinguish between canonical β-catenin-dependent signaling and the two principal non-canonical Wnt branches, planar cell polarity (PCP) signaling and Ca^2+^ signaling. Consistent with the mixed expression of Wnt agonists and antagonists, the strongest downstream enrichment was observed in Ca^2+^ non-canonical Wnt signaling (NFATC1: L_2_FC = 4.10, adj.p = 1.78×10^-163^), with attenuated enrichment of canonical Wnt (LEF1: L_2_FC = 0.63, adj.p = 3.94×10^-9^, **Fig. 3J**). Interestingly, while the Ca^2+^ non-canonical pathway was enriched, the non-canonical PCP pathway appeared comparatively depleted (JUN: L_2_FC = −1.14, adj.p = 1.48×10^-111^), suggesting that plasma-neighborhood-like cells preferentially engage Ca^2+^-mediated non-canonical Wnt signaling, influencing the activation of downstream pathways associated with cancer progression(36).

To determine how alterations in Wnt signaling pathways may influence plasma cell function, we next examined the genes promoted by the enriched transcription factors. Several genes associated with MM cell survival and chemoresistance were expressed at higher levels, including *IRF4*(37) (L_2_FC = 2.00, adj.p = 2.28×10^-196^), *BCL2*(38) (L_2_FC = 1.22, adj.p = 2.76×10^-132^), and *MYC*(39) (L_2_FC = 0.38, adj.p = 1.02×10^-21^, **Fig 3K**).

Furthermore, consistent with promoting CAM-DR, genes downstream of the enriched transcription factors included promoters of homotypic cell-cell adhesion, including *NCAM1*(40) (L_2_FC = 1.87, adj.p = 6.45×10^-91^) and *CDH2*(41) (L_2_FC = 1.83, adj.p = 1.05×10^-28^), potentially supporting MM cell aggregation. Together, these findings suggest that spatially restricted tuning of the Wnt signaling pathways may enhance plasma cell survival, chemoresistance, and cell-cell adhesion through transcriptional programs that reinforce malignant progression and CAM-DR in MM.

### Expression of non-canonical Wnt-promoting agonists is observed across cytogenetic groups and timepoints

Finally, to assess how case-specific factors influence spatial transcriptional programs, we compared plasma-neighborhood-like plasma cells across cytogenetic subgroups and between diagnosis and relapse. Plasma cells from each cytogenetic group were enriched for expected driver genes, such as *FGFR3* in t(4;14) (L_2_FC = 13.07, adj.p = 1.0×10^-250^) and *CCND1* in t(11;14) (L_2_FC = 4.77, adj.p = 1.0×10^-250^, **Supplemental Fig 15A**). Notably, *WNT5B* was expressed at similar levels across patients with or without gain of 1q, loss of 13q, loss of 17p, t(11;14), t(4;14), and hyperdiploidy (all adj.p > 0.05, **Supplemental Fig. 15A-B**) suggesting that non-canonical Wnt signaling represents a conserved feature of plasma cells within plasma cell-predominant neighborhoods, independent of underlying cytogenetics. Consistent with the conservation across cytogenetics, *WNT5B* remained stable at relapse compared to diagnosis, indicating that this signaling axis persists despite therapeutic pressure (all adj.p > 0.05, **Supplemental Fig. 15C-D**). While *WNT5B* expression was consistent across cytogenetics and relapse, other signaling molecules, such as *IL18* in del(17p) samples (L_2_FC = 3.82, adj.p = 3.91×10^-24^) and *IL10* in relapse samples (L_2_FC = 5.87, adj.p = 6.47×10^-24^) were specific to these clinical characteristics, highlighting that additional ligands may further alter the BMME in high-risk and relapsed MM.

### Aberrant ligand expression in plasma-rich neighborhoods associates with immunosuppression and decreased cytotoxicity

After characterizing the spatial expression features of plasma cells in plasma cell-predominant neighborhoods, we next examined how immune cells in these regions are influenced by their local cellular context (**Fig. 4A**). Compared to plasma cells, many non-malignant cell types exhibited far fewer differentially expressed genes (**Fig 4B**). Nonetheless, important biological pathways were represented among the genes that were differentially expressed. Granulocytes and NK cells in plasma cell-predominant neighborhoods had decreased expression of cytotoxic genes including defensins, *DEFA3* (L_2_FC = −1.27, adj.p = 3.25×10^-49^) and *DEFA4* (L_2_FC = −1.17, adj.p = 3.43×10^-53^), as well as the cytotoxic granule protein, *NKG7* (L_2_FC = −1.61, adj.p = 3.55×10^-8^, **Fig. 4C**). Granulocytes additionally had increased expression of *LGALS1*, a known immunosuppressive cell surface protein (L_2_FC = 0.57, adj.p = 2.82×10^-5^)(42). Thus, these observations suggest that even in cell types exhibiting relatively modest spatial transcriptional changes, plasma cell-predominant neighborhoods are associated with reduced cytotoxic gene expression, potentially highlighting a local immunosuppressive microenvironment within the BM.

**Figure 4.**
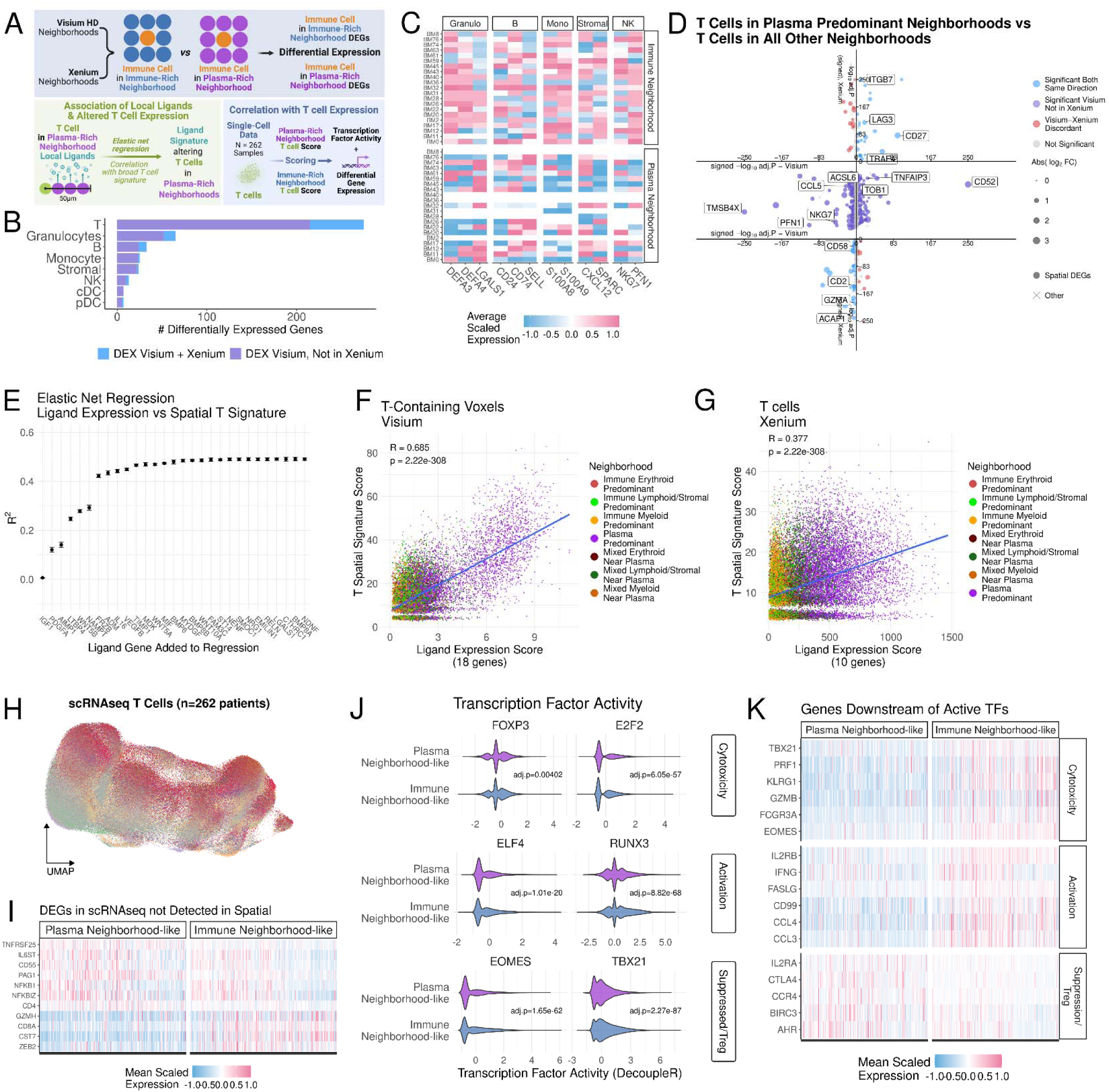
Immune cells in plasma cell-predominant neighborhoods are enrich immunosuppressive genes and decreased cytotoxicity. **(A)** Schematic overview of the spatial analysis framework. Immune cells were compared between plasma cell-predominant neighborhoods and immune-enriched neighborhoods to find spatially differentially expressed genes. Spatially differentially expressed genes were then used to assign spatial labels to single-cell RNA sequenced T cells for higher depth transcriptomic analyses. **(B)** Bar plot displaying the total number of differentially expressed genes by immune cell type. Differential expression was tested using a hurdle model as implemented by the MAST method in Seurat’s FindMakers function with a significance cutoff of multiple comparisons adjusted p value < 0.05. **(C)** Heatmap displaying the mean normalized scaled expression of genes differentially expressed between plasma-neighborhood-like immune cells versus immune-neighborhood-like immune cells in Visium HD data. **(D)** Dot plot displaying the independent differential expression of T cells in plasma cell-predominant neighborhoods versus T cells in immune enriched neighborhoods in both Visium HD and Xenium data. Axes display the -log_10_ adjusted p value signed by the log_2_ fold change (i.e. a gene with a negative log_2_ fold change is displayed on the negative portion of the graph). Genes displaying concordant differential expression (i.e. adjusted p value < 0.05 and log_2_ fold change with the same sign) are displayed in blue, while discordant expression is displayed in red. Given that Visium is a whole-transcriptome assay, many genes in the Visium data were not detected in Xenium data. Genes that were spatially differentially expressed in Visium but not detected in Xenium are in the center of the graph in purple. **(E)** Dot plot showing the association between proximal ligand expression and the spatially differentially expressed genes in T cells. To determine which ligands associated with the observed alterations in T cell expression, elastic net regression was performed between the total expression of each ligand gene within 50 µm of each T cell-containing voxel and the spatial T cell signature comprised of the spatially differentially expressed genes (see Supplemental Figure 16). The ranked ligands were then iteratively added to a combined ligand score to determine the minimum set of ligands that comprise 99% of the association with T cell expression. **(F)** Dot Plot displaying the association between ligand expression and spatial T expression in Visium Data. The X axis shows the combined ligand score of the 18 ligand genes that comprised 99% of the association with T cell spatial expression, and the Y axis displays the spatial T cell expression score. **(G)** Dot plot as in (F) for Xenium data. The X axis displays the ligand score based on the 10 of 18 ligand genes that were detected in Xenium. **(H)** Uniform manifold approximation and projection plot displaying the single-cell RNA sequenced plasma cells colored by patient. See Supplemental Figure 17 for mapping of spatial labels onto single-cell RNA data. **(I)** Heatmap displaying the mean normalized scaled expression of genes that were differentially expressed between plasma-neighborhood-like T cells versus immune-neighborhood-like plasma cells and not well detected in spatial assays. **(J)** Violin plots of transcription activity inferred using DecoupleR. Transcription factors are grouped by row based on the T cell function they promote. **(K)** Heatmap displaying the mean normalized scaled expression of genes that are downstream of the differentially active transcription factors shown in (J).

Of the non-malignant cell types in plasma cell-predominant niches, T cells exhibited the most spatial differential expression, with 305 genes differentially expressed in the V-HD dataset. Of these, 60 were concordantly differentially expressed in the Xenium dataset, 215 were not detected in Xenium, and 40 were detected but either not differentially expressed or differentially expressed in the opposite direction (**Fig 4D**). Similar to NK cells, T cells in plasma cell-predominant neighborhoods displayed reduced cytotoxic potential, with decreased expression of *GZMA* (L_2_FC = −0.82, adj.p = 4.81×10^-3^) and *NKG7* (L_2_FC = −1.59, adj.p = 2.08×10^-47^). Concurrently, T cells also had increased expression of genes known to suppress cytotoxicity, including *TNAIP3*(43) (L_2_FC = 0.67, adj.p = 8.09×10^-26^). Furthermore, T cells in plasma cell-predominant neighborhoods also exhibited signs of chronic antigen stimulation (*CD27*: L_2_FC = 2.31, adj.p = 4.01×10^-93^) and exhaustion (*LAG3*: L_2_FC = 1.18, adj.p = 3.44×10^-19^). Cumulatively, these findings suggest that T cells in plasma-dense regions of the BMME are less cytotoxic, chronically stimulated, and exhausted, potentially reducing their anti-tumoral efficacy.

### Niche-specific ligand expression associates with immunosuppressed T cell expression

To investigate how the signaling milieu of plasma cell-predominant neighborhoods relates to the transcriptional changes observed in T cells, we performed elastic net regression (ENR) comparing ligand expression with T cell gene expression. Briefly, for each T cell-containing voxel, we quantified the total expression of each ligand gene within a 50 µm radius and used these spatial ligand expression values as candidate predictors of the T cell spatial expression score derived from spatially differentially expressed genes (**Fig. 4A**). Ligands were subsequently ranked based on the frequency of inclusion in the ENR model and magnitude of the coefficient between ligand and T cell expression to identify the ligands with the strongest association to altered T cell expression (**Supplemental Fig. 16A**). Then, to identify the minimal set of ligands associated with altered T cell expression, the ranked genes were iteratively added to a growing gene signature. This approach revealed that 18 ligands accounted for 99% of the association between spatial ligand expression and altered T cell expression, suggesting that these ligands collectively contribute to the aberrant transcriptional program observed in T cells within plasma cell-predominant neighborhoods (**Fig. 4E**). Interestingly, a combination of known T-cell-influencing ligands and ligands without well-characterized influences on T cells was found to be associated with altered T cell expression. For example, *MIF* has been reported to promote T exhaustion(23), while *MIF, AIMP1,* and *IGF1* support regulatory T cell (Treg) phenotype (44–46) and *WNT5A* promotes CD8^+^ T cell antigen processing(47), plausibly influencing the observed transcriptional changes in T cells from plasma-rich neighborhoods. Other ligands, such as *TIMP1* and *PDGFA*, are less well-characterized but may represent novel mediators of T cell exhaustion or suppression, potentially acting directly on T cells or indirectly by modulating other cell types within the BMME.

Using the 18 ligand genes to generate a combined ligand expression score, we observed a strong association between proximal ligand expression and the T cell spatial expression score (R = 0.685, p = 2.22×10^-308^, **Fig. 4F**), supporting that T cells in plasma-rich neighborhoods exhibit spatially dependent altered gene expression. Only 10 of the 18 ligand genes were detected in the Xenium data; however, the association of spatial ligand expression and T cell expression was also observed in this orthogonal assay, albeit with a lower R value (R = 0.387, p = 2.22×10^-308^, **Fig 4G**). Cumulatively, these analyses link altered extracellular ligand expression to the transcriptional programs of T cells in plasma cell-predominant neighborhoods, providing evidence that the local signaling milieu may contribute to impaired immune function within these regions.

### Altered transcription factor activity influences T cells in plasma cell-predominant neighborhoods to an immunosuppressive and exhausted state

Given the observed alterations in ligand gene expression within plasma cell-predominant neighborhoods, we next leveraged the greater transcriptomic depth of scRNA-seq to more comprehensively define the associated T cell transcriptional states. As in the plasma cell analysis, we identified T cells in the scRNA-seq dataset that exhibited either plasma-neighborhood-like or immune-neighborhood-like gene expression profiles (**Fig. 3H, Supplemental Fig. 17A-B**). T cells with plasma-neighborhood-like gene expression exhibited enrichment for the expected genes, such as *LAG3* (L_2_FC = 1.57, adj.p = 5.41×10^-95^) and *CD27* (L_2_FC = 1.79, adj.p = 1.0×10^-250^, **Supplemental Fig. 17C**), indicating that the scRNA-seq-defined populations transcriptionally recapitulated T cells identified in spatial assays. Consistent with the reduced expression of cytotoxicity-related genes observed in spatial data, scRNA-seq T cells with plasma-neighborhood-like profiles exhibited an altered CD4/CD8 balance. Specifically, these cells showed increased *CD4* expression (L_2_FC = 0.39, adj.p = 9.51×10^-40^) and decreased *CD8A* expression (L_2_FC −0.58=, adj.p = 3.19×10^-83^) concurrent with increased expression for markers of suppressed T cells such as *CD55*(48) (L_2_FC = 0.27, adj.p = 2.78×10^-48^, **Fig. 3I**). Together, these findings suggest that T cells residing in plasma-rich niches adopt a less cytotoxic and more suppressive transcriptional state.

To define the regulatory programs underpinning this phenotypic shift, we next examined changes in transcription factor activity that could connect plasma-neighborhood signaling cues to T cell functional reprogramming. Notably, several ligands enriched in plasma cell-predominant neighborhoods have been reported to promote transcriptional programs associated with Treg differentiation and function. These include ligands linked to activation of FOXP3, such as *IL16*(49), *AIMP1*(46), *IGF1*(50), and activation of E2F2 by *MIF*(51). Consistent with these observations, T cells with plasma-neighborhood-like expression had increased FOXP3 activity (L_2_FC = 3.35, adj.p = 4.01×10^-3^) and E2F2 activity (L_2_FC = 0.63, adj.p = 6.05×10^-57^, **Fig 3J**). Furthermore, these T cells exhibited reduced activity of critical cytotoxicity-promoting transcription factors, TBX21 (L_2_FC = −0.97, adj.p = 2.27×10^-87^) and EOMES (L_2_FC = −1.20, adj.p = 1.65×10^-62^), further indicating a transcriptional shift towards an immunosuppressed, tumor-promoting BMME.

Having identified shifts in transcription factor activity, we next examined the downstream gene expression programs associated with the differentially active transcription factors. As anticipated, genes linked to Treg function and an immunosuppressive BMME were enriched, including *IL2RA*(52) (L_2_FC = 0.41, adj.p = 1.56×10^-3^) and *AHR*(53) (L_2_FC = 0.44, adj.p = 2.41×10^-33^, **Fig. 3K**). Concurrently, decreased expression of inflammatory and cytotoxic mediators such as *GZMB* (L_2_FC = −0.90, adj.p = 7.13×10^-67^), *PRF1* (L_2_FC = −0.78, adj.p = 6.31×10^-93^), and *IFNG* (L_2_FC = −0.47, adj.p = 6.37×10^-23^) alongside increased expression of the exhaustion-associated marker *CTLA4* (L_2_FC = 0.65, adj.p = 1.50×10^-8^), consistent with a state of T cell exhaustion and depleted effector activity. Together, these findings indicate that plasma-rich niches are associated with a coordinated shift in T cell transcription characterized by enhanced Treg-associated programs and attenuation of cytotoxic effector drivers, supporting the establishment of a suppressed, MM-permissive bone marrow microenvironment.

### LAG3 expression increases at relapse in T plasma-neighborhood-like T cells

Finally, using the scRNA-seq data, we assessed how cytogenetics and disease progression influence the expression of plasma-neighborhood-like T cells. Expression of *LAG3* in plasma-neighborhood-like T cells was stable across patients with/without gain of 1q, loss of 13q, loss of 17p, t(11;14), t(4;14), and hyperdiploid (all adj.p > 0.05, **Supplemental Fig. 18A-B**), indicating that this exhaustion-associated marker is largely independent of underlying cytogenetic alterations. In contrast, *LAG3* was increased at relapse compared to diagnosis (L_2_FC = 0.73, adj.p = 2.0×10^-21^, **Supplemental Fig. 18C-D**), suggesting that the spatially driven marker of T exhaustion is exacerbated with disease progression. Overall, these findings suggest that spatial context plays an important role in shaping T cell transcriptional states, with relapse enhancing features of exhaustion.

### Spatially expressed genes from plasma and immune cells associate with patient outcomes

Having defined cell-type-specific transcriptional profiles using our spatial data, we next evaluated whether these genes were associated with clinical outcomes (**Fig. 5A**). Leveraging our outcome-annotated scRNA-seq dataset (**Fig. 5B**), we partitioned patients into discovery and validation cohorts, consistent with the original publication (8), to assess and validate the prognostic significance of each gene. The expression of each spatially-identified differentially expressed gene was quantified within its corresponding lineage and tested for association with progression-free survival (PFS). Of the 2,054 cell-type-specific differentially expressed genes identified across the Xenium and V-HD datasets, 328 were significantly associated with PFS (p < 0.05) in the discovery cohort, indicating that a substantial subset of spatially-derived differentially expressed genes carries prognostic relevance.

**Figure 5.**
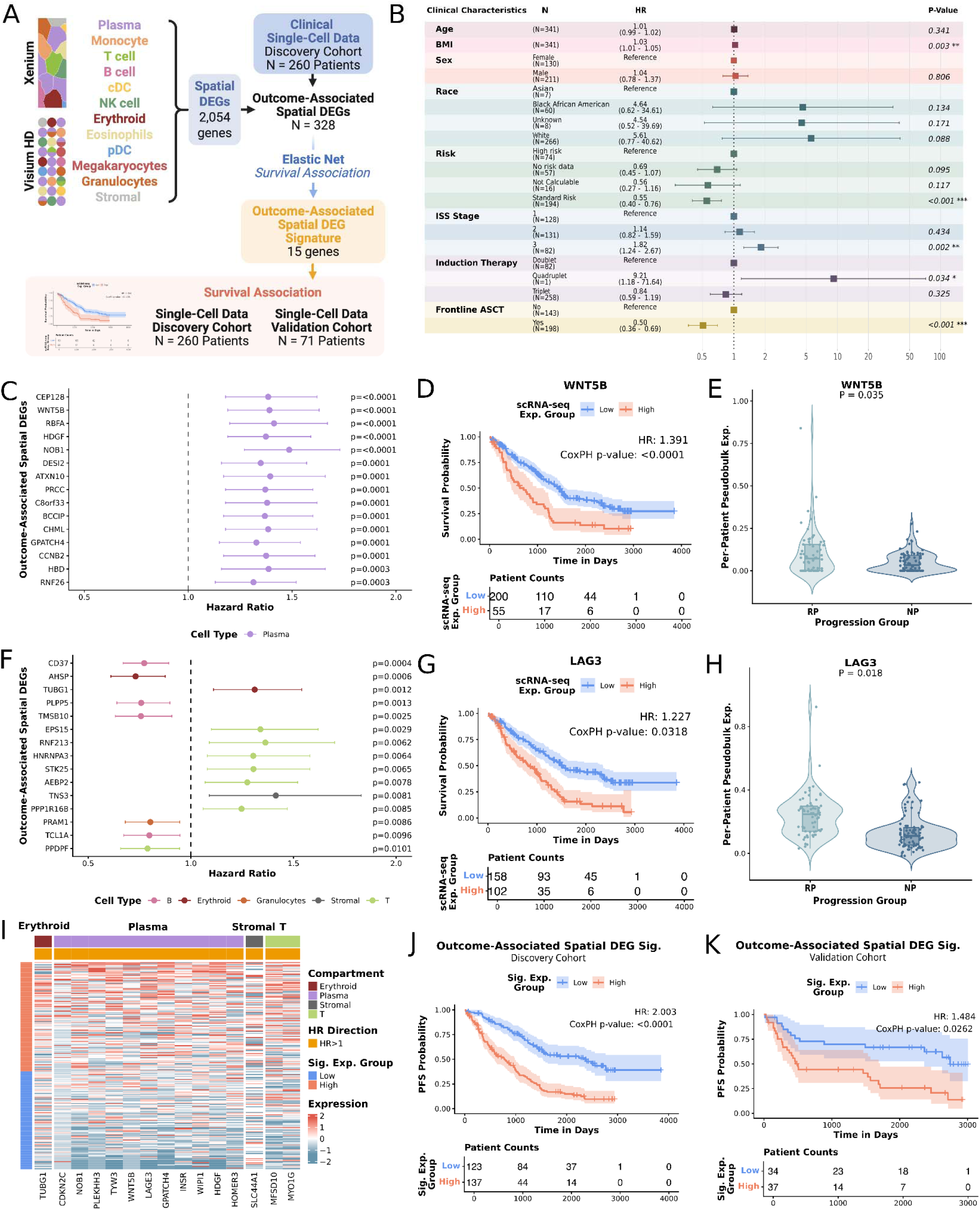
Spatially expressed genes from plasma and immune cells associate with patient outcomes. **(A)** Schematic overview of the analytical workflow. Spatially derived differentially expressed genes (DEGs) were evaluated in the clinically annotated single-cell cohort to identify outcome-associated genes, followed by elastic net regression to derive a 15-gene prognostic signature and independent validation. **(B)** Forest plot showing multivariable Cox proportional hazards model results for clinical covariates in the single-cell cohort and their association with progression-free survival (PFS). Hazard ratios (HRs) and 95% confidence intervals (CIs) are shown. **(C)** Forest plot of the top 15 plasma cell-specific spatial DEGs significantly associated with PFS in the discovery cohort, ranked by Wald test p-value. Points represent HRs, and horizontal lines indicate 95% CIs. **(D)** Kaplan-Meier curves stratified by *WNT5B* per-patient pseudobulk expression in plasma cells, displaying the association with PFS with HR and Wald test p-value. **(E)** Violin plot of per-patient plasma cell *WNT5B* pseudobulk expression in rapid progressors (RP; progression within 18 months) and non-progressors (NP; remission >4 years). Two-sided p-value shown. **(F)** Forest plot of the top 15 spatial DEGs from immune compartments significantly associated with PFS, ranked by Wald test p-value. **(G)** Kaplan-Meier curves stratified by T cell *LAG3* per-patient pseudobulk expression, displaying the association with PFS with HR and Wald test p-value. **(H)** Violin plot of per-patient T cell *LAG3* pseudobulk expression in RP versus NP patients. Two-sided p-value shown. **(I)** Heatmap of scaled per-patient pseudobulk expression of the 15-gene outcome-associated spatial DEG signature across plasma and immune compartments in the discovery cohort. Patients are ordered by composite signature score and stratified into high and low groups as defined in **(J)**. **(J-K)** Kaplan-Meier curves showing the association between the 15-gene composite signature score and PFS in the single-cell discovery cohort **(J)** and independent validation cohort **(K).**

Notably, among the top fifteen outcome-associated spatially differentially expressed gene in plasma cells (**Fig. 5C**), *WNT5B* displayed the strongest association with PFS, with patients with higher plasma cell *WNT5B* expression exhibiting significantly shorter PFS (HR = 1.391, CoxPH p = 4.4×10^-05^, **Fig. 5D**). Consistent with this finding, *WNT5B* expression was also significantly elevated in rapid progressors (RP; progression within 18 months) compared to non-progressors (NP; remission >4 years) (adj.p = 0.035; **Fig. 5E**). This aligns with our earlier spatial observation that plasma cells in plasma-dense neighborhoods are enriched for non-canonical Wnt signaling (**Fig. 3C-D**), further linking this spatially restricted ligand expression to poor clinical outcomes.

Across immune compartments, 58 spatially differentially expressed genes were significantly associated with PFS (p < 0.05; **Fig. 5F, Supplemental Table 1**). Interestingly, among these, increased *LAG3* expression in T cells was associated with inferior PFS (HR = 1.227, CoxPH p = 0.0318; **Fig. 5G**). Furthermore, significantly elevated *LAG3* expression was also observed across T cells in RP compared to NP patients (adj.p = 0.018; **Fig. 5H**). Spatially, *LAG3* expression was upregulated in T cells within plasma-rich neighborhoods (**Fig. 4J**), providing concordance between spatial localization and outcome association. Given that *LAG3* encodes an inhibitory receptor linked to T cell suppression, these findings support that altered transcriptional profiles of T cells within plasma-dense niches are correlated with adverse clinical outcomes.

Next, to refine these results into a prioritized list of prognostic genes, we applied ENR to the 328 survival-associated spatially differentially expressed genes to yield a 15-gene signature spanning plasma and immune lineages (**Fig. 5I**). In the discovery cohort (N = 260), higher composite expression of the resulting outcome-associated gene signature was strongly associated with inferior PFS (HR = 2.003, CoxPH p = 4.4×10^-05^; **Fig. 5J**). Importantly, this association was preserved in an independent validation cohort (N = 71), where elevated signature expression remained significantly associated with shorter PFS (HR = 1.484, CoxPH p = 0.0262; **Fig. 5K**), supporting robustness of the signature.

Collectively, these results demonstrate that transcriptional programs enriched in spatially defined plasma-rich neighborhoods are linked to clinical progression. Specifically, these plasma-rich niches are characterized by plasma cell-mediated noncanonical Wnt signaling associated with early progression. Concurrently, T cells within these niches exhibit altered expression profiles, including increased expression of the inhibitory receptor *LAG3,* which was associated with early progression, consistent with impaired cytotoxic function and immunosuppressive signaling. The combined 15-gene spatial DEG signature integrates these malignant and immune features and stratifies patients by progression risk across independent cohorts, underscoring the prognostic relevance of spatially derived transcriptional profiles.

### Extramedullary MM exhibits spatial enrichment of non-canonical Wnt signaling and T cell suppression

Lastly, to assess how spatial gene expression in extramedullary (EM) lesions compares to the BMME, we performed V-HD on biopsies from five EM lesions (**Fig. 6A**). Samples were collected from EM lesions in the perihepatic soft tissue (n = 1), lymph nodes (n = 3), and periarticular soft tissue (n = 1), and processed using the same quality control and deconvolution procedures as the BM samples (**Fig. 6B, Supplemental Fig. 19A-H**). As expected, >90% of voxels in all five EM samples were annotated as plasma cells (**Fig. 6C-D**). Erythroid cells, megakaryocytes, and eosinophils were not detected, but the remaining non-malignant cell types exhibited canonical gene signature enrichment in EM lesions comparable to that observed in BM samples, indicating that the analytical pipeline performed reliably in this distinct context (**Fig. 6E**).

**Figure 6.**
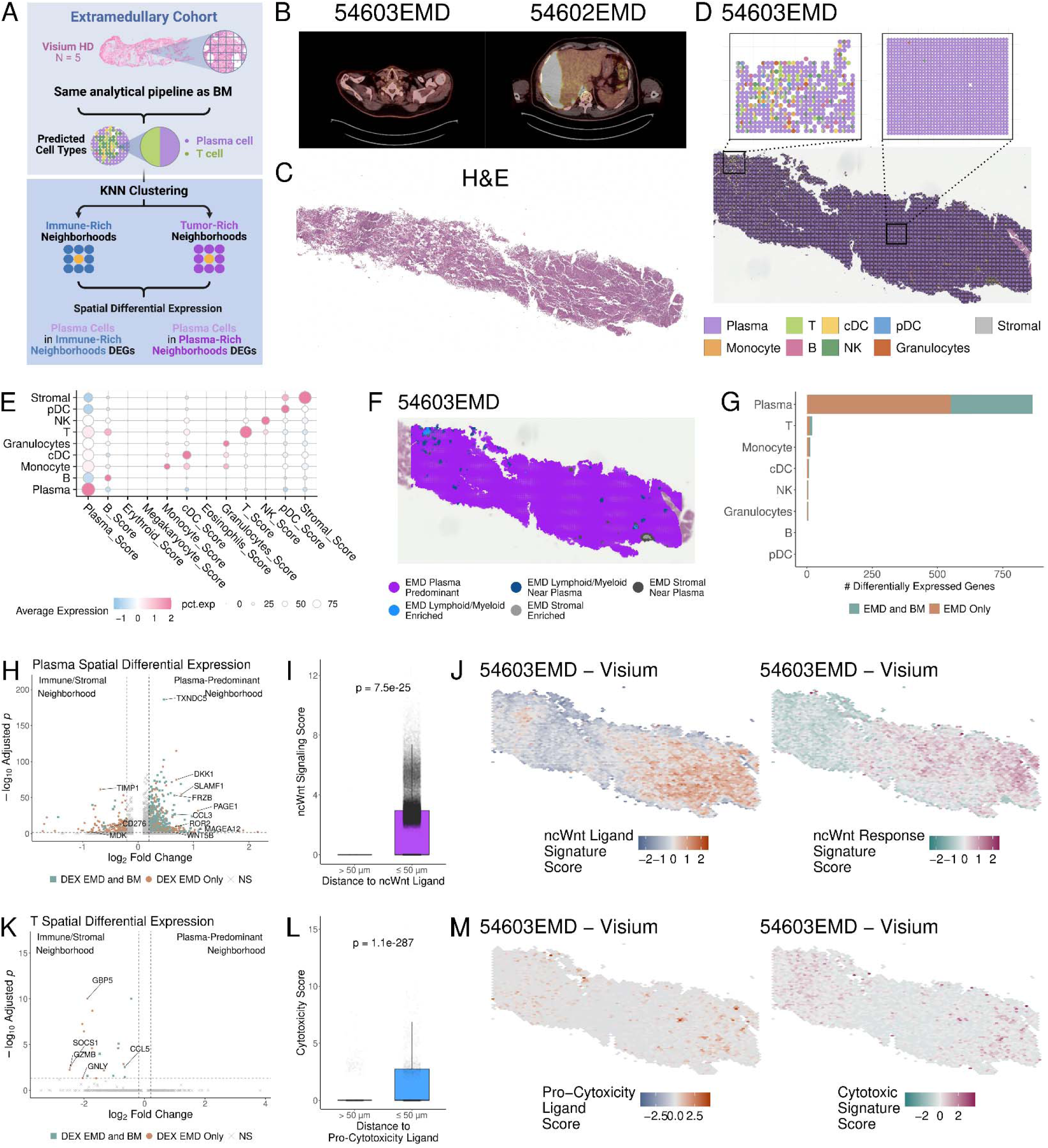
Extramedullary lesions exhibit further enrichment for non-canonical Wnt signaling. **(A)** Schematic overview of the analytical framework for extramedullary (EM) myeloma samples. EM biopsies were analyzed using the same deconvolution, annotation, and clustering pipelines as was used for bone marrow. **(B)** Representative positron emission tomography images of the extramedullary lesions for 54602EMD and 54603EMD. **(C)** Hematoxylin and eosin staining of the EM biopsy of 54603EMD. **(D)** Representative spatial scatterpie plot showing the annotation of voxels in 54603EMD. Voxels are colored by cell type, when multiple cell types are present in a given voxel the scatterpie plot is equally split and colored by the cell types present. **(E)** Dot plot displaying the lineage marker gene scores for the cell types detected in the EM samples. Lineage marker gene scores were calculated as the scaled average normalized expression across 8-12 lineage marker genes per cell type (see Methods for gene lists). **(F)** Representative spatial dot plot displaying the neighborhood annotations for 54603EMD after unsupervised K-nearest neighbor neighborhood analysis. See Supplemental Figure 20 for plots supporting the neighborhood process. **(G)** Bar plot displaying the total number of spatially differentially expressed genes by cell type. For each cell type, differential expression was executed between cells in plasma cell-predominant neighborhoods versus immune/stromal-enriched neighborhoods. Differential expression was tested using a hurdle model as implemented by the MAST method in Seurat’s FindMakers function with a significance cutoff of multiple comparisons adjusted p value < 0.05. **(H)** Volcano plot showing the results of differential expression between plasma cells in plasma cell-predominant neighborhoods versus plasma cells in immune/stromal-enriched neighborhoods. Dots are colored by whether the gene was also differentially expressed in the same comparison for the bone marrow (green), only in the EM comparison (orange), or neither (gray). **(I)** Box plot displaying the plasma cell expression of genes downstream of non-canonical Wnt signaling (Gene ontology pathways GO:2000052 and GO:00900090 as in Figure 4) based on proximity to the nearest voxel expressing a non-canonical Wnt ligand. P value calculated by t-test. **(J)** Representative spatial hexbin plots showing the localization of non-canonical Wnt ligand expression (left panel) and downstream signaling scores (right panel). Color scale represents relative density of ligand expression (blue-white-orange) and relative density of downstream signaling expression (green-white-pink). **(K)** Volcano plot showing the results of differential expression between T cells in plasma cell-predominant neighborhoods versus T cells in immune/stromal-enriched neighborhoods as in H. **(L)** Box plot displaying the T cell expression of cytotoxicity based on proximity to the nearest voxel expressing a cytotoxicity-promoting ligand (see Methods for gene lists). P value calculated by t-test. **(M)** Representative spatial hexbin plots showing the localization of cytotoxicity-promoting ligand expression (left panel) and downstream cytotoxic gene signature scores (right panel). Color scale represents relative density of ligand expression (blue-white-orange) and relative density of downstream signaling expression (green-white-pink).

Consistent with the predominance of plasma cells, the majority of each EM biopsy was classified as plasma cell-predominant neighborhoods (**Fig. 6F, Supplemental Fig. 20A-E**). Small immune and stromal neighborhoods were detected, though they did not contain the compositional diversity observed in the BM; accordingly, neighborhoods were grouped as “lymphoid/myeloid” or “stromal” to reflect the observed composition (**Supplemental Fig. 20A-E**).

To characterize the influence of neighborhood composition on gene expression within EM lesions, we performed spatial differential expression analysis for each cell type across neighborhoods using the same approach as before. Plasma cells exhibited the largest number of spatially differentially expressed genes (n = 864; **Fig. 6G**), likely due to being the most numerous cell type. Of these, 313 genes were concordantly differentially expressed in BM plasma-rich neighborhoods, while 551 were unique to EM lesions. Notably, the non-canonical Wnt ligand, *WNT5B* (L_2_FC = 0.62, adj.p = 8.58×10^-3^) and the canonical Wnt antagonist, *FRZB* (L_2_FC = 0.65, adj.p = 6.44×10^-54^) were enriched in plasma cell-predominant neighborhoods, as was observed in the BM. Furthermore, the canonical Wnt antagonist, *DKK1* (L_2_FC = 0.69, adj.p = 1.73×10^-75^), and the non-canonical Wnt receptor, *ROR2* (L_2_FC = 0.61, adj.p = 4.92×10^-9^) were found to be enriched in EM lesions but were not enriched in the BM (**Fig. 6H**), suggesting that Wnt signaling may become increasingly skewed toward non-canonical pathways in advanced disease.

### Extramedullary MM recapitulates the spatial plausibility of non-canonical Wnt signaling in plasma cell-predominant neighborhoods

To evaluate the spatial plausibility of enhanced non-canonical Wnt signaling, we performed proximity analysis with the same approach as in the BM, quantifying downstream non-canonical Wnt target gene expression relative to distance from non-canonical Wnt-skewing ligands. Consistent with the BM, plasma cells near ligand-expressing voxels exhibited higher downstream target gene expression, supporting a spatially localized influence of these ligands on plasma cell transcriptional programs (**Fig. 6I-J**). Collectively, these results indicate that EM plasma-rich neighborhoods exhibit comparable spatial Wnt signaling to the BM, highlighting a potentially critical neighborhood-dependent shift in pathway activity associated with extramedullary disease.

Finally, to assess the influence of neighborhood composition on T cell transcriptional programs, we performed spatial differential expression analysis across EM neighborhoods. T cells in plasma-rich neighborhoods displayed a clear depletion of cytotoxic programs, including reduced expression of *GZMB* (L_2_FC = −2.48, adj.p = 5.69×10^-3^) and *GNLY* (L_2_FC = −2.05, adj.p = 4.35×10^-2^, **Fig. 6K**). In parallel, plasma-rich neighborhoods were concurrently depleted for several cytotoxicity-supporting cytokines and chemokines, including *CCL5* (L_2_FC = - 1.68, adj.p = 4.76×10^-44^), *CXCL9* (L_2_FC = −2.09, adj.p = 2.06×10^-29^), and *CXCL10* (L_2_FC = −2.61, adj.p = 2.64×10^-51^, **Supplemental Fig. 21A-E**). Spatial proximity analysis further revealed that T cells located near cells expressing these factors exhibited higher cytotoxic gene expression, whereas such spatial arrangements were infrequent within plasma-rich neighborhoods (**Fig. 6L-M**). Together, these findings suggest that plasma-rich EM niches limit cytotoxic T cell activation, reinforcing a locally immunosuppressive microenvironment that may facilitate MM persistence and progression.

## Discussion

A central goal of the spatial transcriptomic study of MM is the quantification of RNA with sufficient spatial resolution and gene capture to characterize single cells and perform downstream transcriptomic analyses such as differential expression. Recent publications, such as Yip et al.(16), using the fluorescence-based Xenium platform have demonstrated that it is feasible to profile the detailed cellular composition of the BMME, but as the authors point out, in BM tissues, Xenium has not consistently captured sufficient transcriptomic complexity for downstream analyses. Herein, we demonstrate that the sequencing-based V-HD assay yields the inverse of these strengths and challenges: 16 µm voxels capture sufficient transcript counts and diversity for downstream analyses, but the voxel size and grid-based organization can result in the capture of transcripts from multiple cells, limiting true single-cell analysis. Furthermore, neither assay captures the feature depth and complexity comparable to scRNA-seq. Thus, we present an approach that leverages the strengths of each assay: V-HD for whole-transcriptome spatial transcriptomic discovery, Xenium for orthogonal validation at cellular resolution on a subset of the transcriptome, and scRNA-seq for extending spatially-derived findings to a larger cohort with greater sequence depth and coverage. Utilizing this paradigm, we successfully characterized niche-specific expression patterns that influence the function of malignant and non-malignant cells within the MM BMME. We speculate that ongoing refinements of these assays may further enhance the ability to characterize the composition and expression of cells in the BMME. For example, Visium HD studies may benefit from the depletion of over-represented transcripts, and Xenium studies may benefit from utilizing a focused panel of hundreds of genes, as opposed to the typical Xenium 5k panel. Nonetheless, we believe our analytical paradigm maximizes the strengths and compensates for the limitations of the respective platforms, enabling high-resolution characterization of spatially resolved gene expression patterns in individual cells.

A known challenge in sequencing-based spatial transcriptomics is the pervasive detection of highly abundant transcripts, particularly immunoglobulin heavy and light chain genes. Even in bone marrow samples from patients without MM, where plasma cells are expected to be relatively rare, immunoglobulin transcripts can be detected in >90% of voxels(15,18). Despite the improved resolution of V-HD, we observed a similar phenomenon; patients with <20% plasma cell burden still had >90% of voxels positive for immunoglobulin genes. This widespread signal likely reflects a combination of cellular co-occupancy within the 16 x 16 µm voxels comprising the grid-based architecture and ambient RNA contamination, including lateral diffusion of transcripts following cell lysis. Such signal inflation complicates accurate cell-type assignment and risks overestimating the abundance of cells producing over-represented genes, contaminating downstream transcriptional analysis. To address this limitation, we extended existing deconvolution frameworks to enable more robust voxel annotation. Specifically, we integrated RCTD-driven deconvolution(17) with a mixture modeling and empirical Bayesian shrinkage framework designed to stabilize cell-type assignments and account for the disproportionate influence of overly abundant genes. By leveraging broader transcriptional signatures rather than relying on lineage-defining transcripts alone, this approach improved the specificity of voxel annotations. Although cross-detection of ubiquitous lineage genes persisted, annotated cell types demonstrated clear enrichment for canonical markers, such as *CD3E* in T cells, supporting the biological validity of the annotations. Importantly, orthogonal validation in Xenium data and concordant findings in matched scRNA-seq datasets further reinforced the robustness of this strategy. Together, these methodological refinements enabled confident within-cell-type, spatially resolved, transcriptional analyses within the BMME.

Spatial interrogation of plasma cell expression revealed many genes preferentially enriched in neighborhoods predominantly comprised of MM cells. Of these, alterations in Wnt signaling showed the most consistent association between enriched expression of ligands and concordant downstream changes in gene expression and inferred transcription factor activity. Wnt pathway aberrations in MM have generally been found to support MM cell survival and proliferation(54–57), CAM-DR(29,58,59), and the development of osteolytic disease(31,32,60). Our findings extend the existing literature, uncovering that Wnt signaling is predominantly enriched in plasma-dense regions of the BMME and skewed towards the non-canonical Wnt pathway. Furthermore, downstream inferred transcription factor activity, such as that of NFATC1, indicates that MM cells in plasma cell-predominant neighborhoods preferentially utilize the Ca^2+^ non-canonical Wnt pathway in addition to some engagement of the canonical β-catenin pathway. Beyond enrichment in plasma cell-predominant neighborhoods, Wnt ligand expression appears to be a high-risk feature based on the observed association with shortened PFS, higher expression in patients who progress rapidly, persistent or increased expression in samples from disease relapse, and concurrent presence in EM lesions. While the most consistently altered gene across these comparisons was *WNT5B*, it seems probable that multiple ligands and/or receptors contribute to the promotion of malignant features in MM cells. The previously observed poor efficacy of DKK1-targeting immunotherapy in patients with smoldering MM may reflect the fact that MM cells utilize several components of Wnt signaling(61). Future studies delineating the consequences of altered expression of individual Wnt ligands and receptors will be critical to understanding the orchestration of aberrant Wnt signaling in MM, potentially providing new therapeutic targets.

Beyond the aberrations in spatial signaling in plasma cells, immune cells in plasma cell-predominant neighborhoods exhibited several dysfunctional features. Across immune cell types, altered gene expression was indicative of a suppressed transcriptional state with decreased anti-MM potential. T cells, in particular, displayed the greatest changes in spatial gene expression. While the association of T cell features with outcomes in MM can vary across disease and treatment contexts, it is generally observed that decreased cytotoxic/effector T cells and increased regulatory T cells associate with poor prognosis(62–65). Our findings support these findings, identifying that these features are preferentially observed in plasma cell-predominant neighborhoods. Furthermore, we associated altered cytokine expression with transcription factor activity and aberrant downstream gene expression, providing a plausible link between spatial expression and poor-outcome-associated immune phenotypes. Intriguingly, several spatially differentially expressed genes within the immune compartment were specifically associated with outcomes. Most notably, increased expression of the inhibitory receptor *LAG3* in T cells was significantly associated with inferior PFS, consistent with previous studies linking inhibitory receptor signaling to immune evasion and adverse outcomes in MM. Furthermore, integrating outcome-associated spatially differentially expressed genes from both immune and plasma cell compartments, we derived a refined 15-gene signature that remained significantly associated with PFS across independent discovery and validation cohorts. Collectively, these results support the prognostic relevance of spatially derived transcriptional programs and indicate that microenvironmental context captured by spatial transcriptomics can identify gene expression states associated with disease progression.

While this study provides novel insights into the spatial dynamics of MM and the BMME, several limitations exist. First, the size of our cohort precluded spatial sub-analyses across features such as cytogenetics and time point. We utilized the neighborhood-like cells in scRNA-seq to provide a surrogate analysis across these covariates; however, future spatial transcriptomic studies with larger cohort sizes will be invaluable for direct comparisons. Second, there are multiple limitations related to the current sequencing depth of spatial transcriptomic assays. A broad goal of spatial analyses is to profile ligands and receptors in juxtaposed cells. Such expression patterns were rarely observed in these data, likely due to data sparsity, leading us to use neighborhood-based comparisons of ligands and downstream signaling pathways to leverage a greater sample of voxels/cells and genes to infer probable interactions. Additionally, the limited feature capture restricted our ability to delineate cellular subtypes, such as naïve, effector, and memory T cells. As spatial assays are further refined to improve sequencing depth, resolving the spatial interactions between juxtaposed cell types with granular annotations may refine our findings with additional important interactions that shape the MM BMME. Finally, our atlas utilizes non-matched samples across V-HD, Xenium, and scRNA-seq. While this may support the robustness of the findings, it does not allow us to directly account for technical variability across platforms.

Our multimodal spatial transcriptomic approach integrates V-HD, Xenium, and scRNA-seq to overcome the individual limitations of each platform, enabling high-resolution, cell type-specific, and niche-resolved analyses of the MM BMME. This strategy revealed spatially organized aberrations in plasma cell signaling, including preferential enrichment of non-canonical Wnt pathway activity in plasma-dense neighborhoods that associate with disease progression and poor outcomes. Furthermore, an altered signaling ligand repertoire corresponded with dysfunctional immune phenotypes. While technical limitations such as voxel resolution, transcript sparsity, and cohort size remain, our findings demonstrate the feasibility and value of combining sequencing- and fluorescence-based spatial assays with single-cell data to dissect the spatial architecture and transcriptional programs driving MM progression. Collectively, this work provides a framework for future studies aimed at mapping cellular interactions, signaling networks, and potential therapeutic vulnerabilities within the MM microenvironment.

## Methods

### Ethics Approval and participant consent

Informed consent for BM and EMD biospecimen collection and processing in accordance with the Declaration of Helsinki was obtained from all patients under protocols approved by the Emory School of Medicine Institutional Review Board (IRB Study ID 00057236 and 00052278) and Icahn School of Medicine at Mount Sinai Institutional Review Board, including the Multiple Myeloma Biorepository (IRB Study ID 1800456).

### Data availability

The raw and processed data generated in this study have been deposited in the National Center for Biotechnology Information Gene Expression Omnibus and will be made public with publication of this article. The Xenium dataset used in this study is available at GSE299207. The reference scRNA-seq dataset used for voxel deconvolution was constructed from GSE161801, GSE253355, and E-MTAB-14010. The scRNA-seq dataset used to extrapolate spatial findings and correlate clinical outcomes is available under controlled access at MMRF’s VLAB shared resource (https://mmrfvirtuallab.org). The MMRF requires the minimum qualifications for access: apply for access at https://mmrfvirtuallab.org and to meet the following minimum qualifications: (i) must be a permanent employee of their institution and at a level equivalent to a tenure-track professor, (ii) senior investigator that is overseeing laboratory or research program. If the access request is approved, usually within a week, investigators will receive an email with instructions for downloading the data. Alternatively, a Seurat R object with processed UMI counts and limited metadata can be accessed at Zenodo (10.5281/zenodo.15025566).

### Materials Availability

This study did not generate new unique reagents.

### Bone marrow biopsy sample collection

Bone marrow core biopsies were obtained from patients diagnosed with Multiple Myeloma as part of routine clinical evaluation. Biopsy specimens were fixed in neutral buffered formalin, decalcified to enable sectioning of mineralized bone tissue, and subsequently embedded in paraffin to generate formalin-fixed paraffin-embedded (FFPE) blocks.

### Visium HD sample preparation, library construction, and sequencing

Spatial transcriptomic profiling was performed using 10X Genomics V-HD Spatial Gene Expression for FFPE platform according to manufacturers’ instructions. FFPE tissue blocks were sectioned with 5 µm thickness onto VWR Superfrost microscopic slides. The sections were deparaffinized and decrosslinked. Human whole transcriptome probe panels consisting of specific probes for each targeted gene were then added to the tissue. These probe pairs hybridize to their gene target and ligate to one another. Tissue slides and Visium CytAssist Spatial Gene Expression Slides were loaded into the Visium CytAssist instrument, where they were brought into proximity with one another. Each Visium CytAssist HD Slide contains about 11 million Capture Areas (2×2 um) with barcoded spots that include oligonucleotides required to capture gene expression probes. Gene expression probes were released from the tissue upon CytAssist Enabled RNA Digestion & Tissue Removal, enabling capture by the spatially barcoded oligonucleotides present on the Visium slide surface. All the probes captured on a specific spot share a common Spatial Barcode. Libraries were subsequently generated from the probes and sequenced, and the Spatial Barcodes were used to associate the reads back to the tissue section images for spatial mapping of gene expression. Sequencing was performed on an Illumina NovaSeq 2000 sequencer.

### Visium HD dataset processing and quality control

V-HD sequencing data were processed using Space Ranger (v3.0.1, 10x Genomics Inc.) to demultiplex sequencing reads into FASTQ files, align reads to the human reference genome (GRCh38 2020-A), and generate gene-by-voxel UMI count matrices. V-HD uses spatially barcoded oligonucleotide probes to assign captured transcripts to 2 µm x 2 µm capture areas. Adjacent capture areas can be aggregated into larger voxels to provide sufficient transcriptomic depth for downstream analysis. We tested 2 µm x 2 µm, 8 µm x 8 µm, and 16 µm x 16 µm voxels with quality control (QC) thresholds of at least 100 unique molecular identifiers (UMIs) and less than 20% of UMIs mapping to mitochondrial genes. Two µm x 2 µm and 8 µm x 8 µm voxels had insufficient voxels passing QC due to low gene count. We therefore used the 16 µm x 16 µm voxels to maximize the number of QC-passing voxels while retaining near-single-cell spatial resolution.

### Single-cell clinical dataset source, processing, and quality control

Single-cell RNA-sequencing data with clinical annotations were accessed via the MMRF VLAB shared resource. Dataset generation, pre-processing, and the initial annotation were previously reported by Pilcher *et al.*(8) The full dataset comprises 480 bone marrow aspirate samples collected from 347 multiple myeloma patients enrolled in the MMRF CoMMpass study (NCT01454297) at various clinical time points. For downstream analyses, we restricted the dataset to samples collected at baseline (N = 320) and relapse/progression (N = 81).

### Xenium dataset source and quality control

Xenium data from 10 MM bone marrow biopsies produced by Yip *et al.*(16) was accessed via GSE299207. The raw feature barcode matrices were subset to the samples corresponding to MM samples; precursor samples and healthy controls were filtered out. Consistent with Yip *et al.,* Low-quality cells were filtered out if they contained fewer than 50 unique genes, or fewer than 60 total counts.

### Voxel annotation overview

Voxels in V-HD data frequently capture transcripts from multiple cells. This arises from several factors, including imperfect alignment between V-HD grid-based capture areas and the *in-situ* arrangement of cells, the voxel size (16 µm x 16 µm) compared with many bone marrow cell types, and lateral diffusion of transcripts following cell lysis. Cell type identification is further complicated by the widespread detection of highly abundant transcripts. For example, immunoglobulin heavy and light chain transcripts are often over-represented in Visium bone marrow datasets, including in samples from healthy control patients where plasma cells are expected to be rare (18). Consistent with this phenomenon, we observed that in most samples >90% of voxels contained transcripts from a patient’s involved immunoglobulin heavy or light chain gene, even when the histologically quantified plasma cell percentage was <20%.

To identify the primary cell type(s) contributing to the transcriptomic profile of each voxel while accounting for the intrinsic imbalance in gene representation, we developed a three-part analytical framework. First, voxels were deconvolved using a reference single-cell RNA sequencing dataset to estimate the proportional contribution of each cell type to each voxel. Second, the estimated cell type proportions were fit to a logistic–gamma mixture model to derive a preliminary threshold for binarizing voxels (e.g., “plasma-containing voxel”) based on expected cell composition. Third, empirical Bayesian shrinkage procedure was applied to account for deviations between the observed and expected distributions of cell type signals. Specifically, when the observed distribution of a given cell type differed from that predicted by the mixture model (e.g., excess T cell signal), the threshold was adjusted to better reflect the empirical distribution. Each step of this framework is described in detail in the following sections.

### Construction of reference dataset for voxel deconvolution

During quality control and initial exploration of the V-HD data, we observed clear expression of canonical marker genes for eosinophils and stromal cells. Because these cell types are typically underrepresented in scRNA-seq datasets, we constructed a reference dataset from three complementary sources: (1) an unsorted MM bone marrow biopsy scRNA-seq dataset (GSE161801), (2) stromal cells from a dataset in which stromal populations were enriched prior to sequencing (GSE253355), and (3) a dataset of sorted eosinophils (E-MTAB-14010). All datasets were processed using a consistent analytical pipeline. Low-quality cells were filtered out if they contained fewer than 200 unique genes, greater than 7500 unique genes (presumed doublets), or more than 10% mitochondrial reads. Raw counts were log-normalized using a scaling factor of 10,000 to account for differences in sequencing depth across cells. Highly variable genes were identified, and principal component analysis (PCA) was performed on the top 2,000 variable genes to reduce dimensionality. The first 20 principal components were used for Louvain clustering and visualization with Uniform Manifold Approximation and Projection (UMAP). Clusters were annotated into cell types based on canonical marker gene expression. After quality control and annotation, high-quality stromal cells and eosinophils were merged with the annotated MM whole bone marrow dataset to generate the final reference dataset used for voxel deconvolution.

### Voxel deconvolution

To estimate the proportional contribution of reference cell types to the transcriptional profile of each voxel, deconvolution was performed using the Robust Cell Type Decomposition (RCTD) algorithm implemented in the spacexr R package(17). RCTD estimates the cellular composition of voxels by modeling observed gene expression as a mixture of cell type-specific expression profiles derived from the scRNA-seq reference dataset described in the previous section. RCTD models gene counts in each voxel using a probabilistic framework based on a Poisson distribution, while accounting for differences in sequencing depth and platform-specific effects between the spatial and reference datasets. We ran RCTD in “full” mode, which utilizes maximum likelihood optimization to estimate the proportion of all reference cell types (as opposed to doublet mode which estimates the relative contribution of, at most, the two most represented cell types). This approach allows complex cellular compositions to be inferred within each spatial capture location. The result of this analysis is a voxel by cell type matrix where each entry represents the estimated proportion of that voxels transcriptome coming from each cell type. This matrix was used in the subsequent two steps to estimate cut points for binarizing voxels as “containing” or “not containing” a given cell type (e.g. a plasma cell containing voxel).

### Derivation of cut points for labeling voxels as containing cell type(s)

To create annotation thresholds for each cell type we performed a two-step approach. First, we fit a hurdle model to the observed proportions across voxels for each cell type and sample. The hurdle model was composed of a logistic regression component modeling near-zero detection of a given cell type, and a gamma model for the left-skewed non-zero proportion values. Then, we calculated a preliminary cut point for each cell type based on the expected proportion or each cell type in the sample. For plasma cells, we used the histologically estimated plasma cell proportion. For example, in a sample with 50% plasma cells, the threshold for a “plasma-containing voxel” would be set at the 50^th^ percentile of the plasma cell logistic-gamma hurdle model. For the non-plasma cell types, we derived preliminary estimations from review of scRNA-seq(8,66), CO-Detection by indEXing(66), and mass cytometry (CyTOF)(67,68) datasets to provide expected proportional abundance robust to estimates from any single assay. The expected cell type proportions for the non-malignant compartment were: 0.08 B, 0.03 cDC, 0.02 eosinophils, 0.21 erythroid, 0.22 granulocyte, 0.01 megakaryocyte, 0.09 monocyte, 0.04 NK, 0.01 pDC, 0.08 stromal, and 0.21 T. Thus, for a sample with 50% plasma cells, the preliminary threshold for T cells would set as the 89.5th percentile of the T cell logistic-gamma hurdle model. The result of this procedure was a list of preliminary cut points for each cell type and sample.

Second, we applied an empirical Bayes shrinkage procedure to adjust the preliminary cut points based on the observed distribution of the deconvolution data. For example, if T cell transcriptomic profiles were overrepresented in a given sample relative to the literature-derived expectation, the shrinkage procedure decreased the annotation cut point for T cells in that sample to more accurately reflect the observed gene expression. For a vector of cut points, y, the mean (μ̂) and variance (σ̂^2^) were first computed. The between-sample variance component (î^2^) was then estimated as max(0, σ̂^2^ - σ̂^2^/r), where r denotes the number of observations. A shrinkage weight was subsequently defined as w = τ̂^2^/ τ̂^2^ + σ̂^2^. Each observation was then adjusted toward the global mean using a convex combination 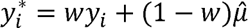. The result of this procedure is to shrink individual values toward the overall mean when the estimated variance is small relative to sampling noise. Effectively this decreases unexpectedly high cut points that are likely to represent samples with more of a given cell type than expected, and increases unexpectedly low cut points that are likely to represent samples with less of a given cell type than expected.

### Optimization of annotation reference granularity

To optimize the granularity with which to annotate cells, we tested several levels of granularity for each cell type in the reference dataset. For example, we tested annotating T/NK lymphoid cells as (i) a single T/NK label, (ii) two labels T and NK, (iii) three labels including CD4^+^ T, CD8^+^ T, and NK. Similar levels of granularity were tested for each cell type. Voxels were then deconvolved and annotated using the approach described in the previous sections. To assess annotation quality, differential expression was performed between voxels containing a given cell type and all other voxels. The optimal granularity was selected based on the highest log_2_ fold-change of canonical cell type genes (e.g. *CD3E* in T cells, *SDC1* in plasma cells, etc.).

### Assessment of annotation quality in Visium HD dataset

To assess the quality of optimized voxel annotations we performed one-versus-all cell type differential expression to evaluate the canonical cell type gene expression. Additionally, we curated cell type gene lists and calculated cell type expression scores from the scaled, normalized counts of all genes in each list. The gene lists for each cell type were as follows: B cells were scored using the genes *TCL1A, CD22, LTB, SPIB, BIRC3, BCL11A, LY86, FOXP1, SNX2, RCSD1, CD24, LIMD2, STX7, RASGRP2,* and *DCK*. Conventional dendritic cells (cDCs) were scored using *CLEC10A, FCER1A, LGALS2, ITGAX, SAMHD1, IGSF6, CXCL10, C1orf162, IFI30, RNASE6, LY86, CTSH, GLIPR1, S100A10, CFP,* and *FGL2*. Eosinophils were scored using *MMP25, PADI2, CEACAM3, CLC, VSTM1, FES, S100P, PYGL,* and *PGLYRP1*. Erythroid cells were scored using *GYPA, KLF1, CA1, HBM, CA2, MYL4, RHCE, ACHE,* and *AHSP*. Granulocytes were scored using *CTSG, PRTN3, ELANE, AZU1, MPO, PRSS57, MS4A3, CLEC11A, DEFA3, DEFA4, DEFB1, CST7, PLPPR3, PRRT4,* and *CEACAM6*. Megakaryocytes were scored using *LY6G6F, TMEM40, ITGA2B, ITGB3, PTCRA, CLEC1B, PF4, PPBP, TREML1, GP6, CMTM5, SELP,* and *GP9*. Monocytes were scored using *S100A12, LILRA5, PLBD1, MNDA, CFP, FOLR3, CRISP3, FGR, CLEC7A, CD14, CAMP, ITGAM, C5AR,* and *SERPINA*1. Natural killer (NK) cells were scored using *GNLY, S1PR5, PRF1, CD247, MATK, IL2RB, GZMA, KLRD1, KLRB1, CD7, CTSW, GZMB,* and *NKG7*.

Plasmacytoid dendritic cells (pDCs) were scored using *IL3RA, TLR7, SPIB, SERPINF1, IRF7, TGFBI, PTGDS, TCF4, IRF8, TPM2,* and *APP*. Plasma cells were scored using *CD38, XBP1, TNFRSF17, SSR4, MYCN, TMEFF1, SLAMF7,* and *MZB1*. Stromal cells were scored using *CHAD, COL5A2, COL5A1, BGLAP, IBSP, CPE, MYH11, PTH1R, ACTA2, SPP1, ELN, TBX2, SPARC, OMD, CREB3L1, LRP4, MMP13,* and *SERPINF1*. T cells were scored using *TRAC, CD3D, TRGC2, CD8A, TRBC2, GZMK, KLRG1, IL32, CD2, TRBC1, CD3E,* and *CCL5*.

### Cell type annotation in Xenium dataset

Cell types in the Xenium dataset were annotated to harmonize with the optimized annotations in the V-HD datasets. As with scRNA-seq data, raw counts were log-normalized using a scaling factor of 10,000 to account for differences in gene detection across cells. Highly variable genes were identified, and PCA was performed on the top 2,000 variable genes to reduce dimensionality. The first 20 principal components were used for Louvain clustering and visualization with UMAP. Clusters were annotated into cell types based on canonical marker gene expression.

At the full bone marrow level, plasma cells, stromal cells, erythroid cells, megakaryocytes, and immune cells were differentiable, however, lymphoid and myeloid cells did not clearly form discrete clusters. To achieve more precise annotation of immune cell types, we repeated the dimension reduction and clustering process for cells in the immune compartment, using variable genes specific to that compartment. This process enabled annotation of the 12 cell types annotated in the V-HD dataset, harmonizing the cell types analyzed.

### Cell type annotation in clinical single-cell RNA sequencing dataset

The scRNA-seq dataset from Pilcher et al.(8) was processed using Seurat (v5.4.0)(69). Using the top 2,000 variable genes, principal component analysis (PCA) was performed, and the first 30 principal components were used for dimensionality reduction. To correct for batch effects, we applied Harmony (v1.2.4)(70) to the PCA-embeddings, specifying each unique combination of processing center and shipment batch as a covariate. Louvain clustering was then performed on the batch-corrected Harmony embeddings using the first 30 principal components and a resolution parameter of 1.0 via Seurat’s ‘FindClusters’ function. Clusters were visualized using UMAP.

To facilitate integration with spatial transcriptomic analyses, cell type annotations were harmonized with those used in the spatial datasets. Original single-cell cluster-level annotations reported by Pilcher *et al.*(8) were consolidated into the broader lineage labels applied to the deconvolution and annotation of the Visium and Xenium datasets (**Supplementary Table 2**).

### Cellular neighborhood derivation in Visium HD and Xenium datasets

To evaluate the influence of local cellular composition on gene expression, we performed unsupervised neighborhood clustering analysis. For each voxel, we quantified the proportion of all cell types within a fixed radius. The result was a voxel-by-cell type matrix where each entry represented the proportion of the corresponding cell type within the radius of the corresponding voxel. We initially tested radii from 16 µm (i.e. one voxel width) to 250 µm representing a conservative maximum plausible distance over which a cell could communicate with the index voxel via a paracrine signaling molecule(71). We found that 50 µm provided the most informative neighborhood definition, producing clusters that appeared enriched for specific cellular compositions. Smaller radii captured too few surrounding cells and therefore did not adequately represent the proximal cellular environment, whereas larger radii increasingly averaged over heterogeneous regions and reduced the specificity of the resulting neighborhoods.

### Differential gene expression analyses

For all comparisons, we utilized the Model-based Analysis of Single-cell Transcriptomics (MAST) method(72). MAST applies a generalized linear hurdle model to account for the high frequency of technical zeroes present in transcriptomic assays such as V-HD, Xenium, and scRNA-seq. The hurdle model is composed of a logistic regression component which models the probability of detecting a gene, and a Gaussian linear model for the log-normalized expression in voxels with non-zero detection. Significance is assessed using a likelihood ratio test comparing the groups specified by each comparison. For example, comparing plasma cell-containing voxels present in plasma cell-predominant neighborhoods versus plasma cell-containing voxels in immune-enriched neighborhoods. P values were adjusted for multiple testing using Bonferroni correction.

Due to the mixed cellular gene profiles captured in some voxels, spurious differential expression can occur when comparing cell types across neighborhoods with differing cellular composition. In such cases, genes identified as differentially expressed may reflect canonical expression of neighboring cell types rather than true transcriptional differences within the cell type of interest. For example, comparing T cells located in plasma cell-predominant neighborhoods to T cells in immune-enriched neighborhoods may yield immunoglobulin genes as differentially expressed due to signal originating from nearby plasma cells. To mitigate these effects, we leveraged our scRNA-seq reference dataset to generate cell type-specific gene lists and filtered spatial differential expression results to retain only genes expressed by the tested cell type. For each cell type, we performed one-versus-all differential expression in the reference scRNA-seq dataset. Genes with a multiple-comparisons-adjusted p-value < 0.05 and log_2_ fold change > 0.2 were included in the gene list for that cell type. In all spatial comparisons, only genes that were present in the corresponding cell type’s gene list were retained.

### Evaluation of expression similarity across neighborhoods in Visium HD and Xenium datasets

To determine which cellular neighborhoods had similar or disparate expression profiles, we performed PCA on the expression of each cell type across neighborhoods. Using the reference cell type gene lists described in *Differential gene expression analysis*, we identified the top 200 highly variable genes for each cell type across neighborhoods. As an example for plasma cells, we subset the V-HD dataset to the genes present in the reference plasma gene list, and to the plasma-containing voxels. Then we identified the top 200 highly variable plasma cell genes across the seven identified neighborhoods for use in PCA. Then, the average Euclidean distance using the top 5 PCs was calculated to provide a standardized metric where lower values indicated similar expression profiles and higher values indicated dissimilar expression profiles. This procedure was repeated across all 12 cell types analyzed in the spatial datasets. The same method was used for V-HD and Xenium datasets.

### Identification of differentially expressed ligands, receptors, and concordant pathway genes

To identify which of the differentially expressed genes coded for ligands and receptors, we used a multi-step process. Candidate ligand and receptor genes were first identified using Gene Ontology (GO) annotations using the biomart package(73). To generate an initial list of candidate ligands, we retrieved genes annotated with GO terms related to ligand activity or extracellular signaling, including GO:0005125, GO:0005109, GO:0008083, GO:0005102, and GO:0005615. For receptors, genes were retrieved based on GO terms associated with receptor-mediated signaling, including GO:0007166, GO:0004872, GO:0004888, and GO:0038023. Unique HGNC gene symbols from these queries were compiled to produce broad candidate lists of ligand and receptor genes.

Given that GO annotations can include genes that participate indirectly in ligand and/or receptor function, candidate lists were further curated to retain genes most likely to encode bona fide secreted ligands or membrane receptors. Ligands and receptor genes were cross referenced against annotations in UniProt (https://www.uniprot.org/) the Human Protein Atlas (https://www.proteinatlas.org/) and ligand-receptor interaction resources including CellPhoneDB(74). Genes lacking evidence for secretion (for ligands) or membrane receptor activity were removed, resulting in a refined set of high-confidence ligand and receptor genes used for downstream analyses.

After identification of differentially expressed ligand genes, we similarly used biomart to identify gene lists that corresponded to pathways downstream of enriched ligands and receptors. The goal of this analysis was to provide evidence for which ligands were influencing expression in plasma cells. For example, we observed that *WNT5B* was highly expressed in plasma cells from plasma cell-predominant neighborhoods. To determine if *WNT5B* was likely influencing plasma cell expression (instead of, or in addition to, influencing other cell types), we utilized the GO pathway GO:2000052 “non-canonical Wnt signaling pathway”. By finding both spatial differential expression of ligands, and spatial differential expression of genes downstream of that ligand, we filtered candidate genes to those with the most evidence for a meaningful impact on signaling and plasma cell transcription.

### Spatial analysis of ligand expression and concordant pathway gene expression

From the neighborhood analyses, we had identified over-expressed ligands with concordant enrichment for pathways downstream of each ligand. To evaluate the spatial plausibility of ligand expression and downstream pathway enrichment, we quantified the expression of downstream pathway genes as a function of distance to the nearest ligand-expressing voxel. Given plasma cell-predominant neighborhoods exhibited enrichment for both non-canonical Wnt agnoists (e.g. *WNT5B*) and canonical Wnt antagonists (e.g. *FRBZ*), concurrent with differential expression of several genes in the GO:2000052 “non-canonical Wnt signaling pathway” and GO:00900090 “negative regulation of canonical Wnt signaling pathway”, we combined these features into a single, non-canonical Wnt promotion analysis. Downstream signaling enrichment was calculated for each voxel from the genes present in GO:00900090 and GO:2000052. Voxels were binarized as either expressing or not expression a relevant ligand based on the expression of either a non-canonical Wnt agonist or a canonical Wnt antagonist (*WNT5B, WNT5A, WNT11, DKK1, SFRP1, SFRP2, FRZB, WIF1, SOST, SOSTDC1*). Finally, the relationship between downstream signaling and distance to the nearest ligand expression voxel was evaluated by subsetting to voxels containing the relevant cel type (e.g. plasma cells in the non-canonical Wnt analysis) and grouping voxels in 16 µm intervals to determine if voxels at closer proximity to ligand expression had higher downstream enrichment scores.

### Mapping spatial labels to single-cell RNA sequencing gene expression profiles

Spatial transcriptomic assays typically capture hundreds of UMIs per cell/voxel. By comparison, scRNA-seq can capture tens of thousands of UMIs per cell. To utilize the increased feature complexity of scRNA-seq data, we mapped spatial labels onto the corresponding cell types using spatially differentially expressed genes. From these gene lists, the AddModuleScore function in Seurat was used to calculate a neighborhood-like score. For example, a plasma cell in plasma neighborhood-like score was calculated from the genes differentially expressed by plasma cells in plasma cell-predominant neighborhoods, and the same was executed for plasma cells in immune-enriched neighborhoods. Then, for each cell a difference threshold of 0.1 was applied, such that cells preferentially expressing genes corresponding to a specific neighborhood were labeled “plasma neighborhood-like” or “immune neighborhood-like”. Cells that did not meet the threshold in either direction were labeled “indeterminate”. Differential expression was performed between neighborhood-like groups as described in *Differential gene expression analyses*.

### Transcription factor activity inference in single-cell RNA sequencing dataset

Transcription factor activity was inferred using the decoupler package(75). This package leverages collecTRI, a curated resource that integrates information from 12 transcriptional regulatory databases and provides TF-target interactions with signed weights (+1 for activation and −1 for repression). For each cell, a univariate linear model was fit in which the normalized expression of a transcription factor’s target genes is regressed against the corresponding interaction weights. The t-statistic of the regression slope is used as the transcription factor activity score, reflecting the degree of concordance between observed target expression and the expected regulatory direction. Positive scores indicate activation-consistent expression patterns, whereas negative scores suggest repression or lack of activity. Differential transcription factor activity was assessed using the same approach as differential gene expression.

### Elastic net regression of ligand expression and altered spatial T cell expression

To associate the abundance of ligand genes with altered spatial T cell expression, we applied elastic net regression using spatially aggregated ligand expression as candidate predictors in the V-HD dataset. For each T-containing voxel, we identified all other voxels present within a 50 µm radius. For each candidate ligand gene, the mean expression across neighboring spots within this radius was computed. The response score was calculated using the genes with positive log_2_ fold change values so that score captures the enriched genes from T cells in plasma cell-predominant neighborhoods. Predictor variables in the elastic net regression therefore consisted of the proximal expression of candidate ligand genes, while the response variable was the T cell gene score.

Elastic net regression models were fit using the glmnet package(76) with mixing parameter α = 0.5 to balance L1 and L2 regularization. The regularization parameter for each model was selected with cross-validation by minimizing the mean error. To robustly identify association between ligand expression and altered T expression, we performed 100 bootstrapped iterations, refitting the cross-validated elastic net model on each resampled dataset. The strength of association between individual ligands altered T cell expression was calculated from the median regression coefficient across bootstraps multiplied by the frequency of ligand gene inclusion in the optimal elastic net model.

Then, to identify the minimum set of ligand genes that explained 99% of the association between ligand expression and altered T cell expression, we performed iterative feature addition. Ligand genes were ranked based on their association strength and iteratively added as predictor variables in a regression model against the same spatial T score as used in elastic net regression. On each iteration we fit a regression model with the ligand genes as predictors using the caret package(77). Model performance was assessed using 10-fold cross-validation, and predictive accuracy was quantified by the cross-validated coefficient of determination (R^2^). Finally, to identify the minimum ligand gene list that accounts for 99% of the association between ligand expression and altered T cell expression, we calculated the number of genes required to achieve 99% of the maximum observed R^2^ value across all ligands. The resulting genes were considered the ligands most likely to be contributing to the observed alteration in T cell expression.

The Xenium dataset was used as orthogonal validation of the V-HD-derived association between ligand abundance and T cell expression. Because Xenium is not a whole-transcriptome assay, some of the ligands in V-HD were not present in Xenium. We therefore utilized the subset of the minimum ligand gene set that were present in both assays. For each ligand gene, the expression within a 50 µm radius of each T cell was calculated as described above. The T response score was calculated using the subset of the spatially differentially expressed genes list that was also present in Xenium data. As with the V-HD analysis, we fit a regression model with the ligand genes as predictors of T expression and model performance was assessed using 10-fold cross-validation.

### Survival Analyses

For survival analyses, patients with available outcome data were partitioned into discovery (N = 262) and validation cohorts (N = 77), as reported by Pilcher *et al*. To evaluate the association between spatially derived differentially expressed genes (DEGs, N = 2,054) and progression-free survival (PFS), cell type specific psuedobulk expression profiles were generated for each patient in the discovery cohort. Briefly, the single-cell object was subset to the respective cell type, followed by the aggregation of gene counts for each patient, log-normalization, and scaling using the Seurat ‘PseudobulkExpression’ function.

Associations between gene expression and PFS were assessed using Cox proportional hazards regression implemented with the survival package (v3.8.3). For each gene, scaled pseudobulk expression was modeled as a continuous variable with age, body mass index (BMI), receipt of frontline autologous stem cell transplantation (ASCT), and the coupled batch number and study site (site-batch) from sample processing as covariates. Hazard ratios (HRs) with 95% confidence intervals (CIs) and two-sided Wald test p-values were extracted from model coefficients. For visualization, patients were stratified into high- and low-expression groups based on the optimal cut point value calculated using the survMisc package (v0.5.6). Kaplan-Meier survival curves were generated using the survminer package (v0.5.1).

### Outcome-associated spatial gene signature

To derive a prognostic signature from outcome-associated spatially differentially expressed genes (DEGs), penalized Cox proportional hazards regression with elastic net regularization was performed using the glmnet package (v4.1.10) (76,78). Modeling was restricted to patients in the discovery cohort with available PFS data (N = 260). For each gene, cell-type-specific pseudobulk expression values were calculated per patient. Elastic net models were fit using the scaled pseudobulk expression values for all candidate DEGs as predictors and PFS as the outcome. To identify robust prognostic features, the elastic net mixing parameter (α) was evaluated across a range of values from 0 to 1 in increments of 0.1. For each α value, 10-fold cross-validation was used to determine the optimal regularization parameter (λ) with the minimum cross-validated partial likelihood deviance. Genes with positive non-zero coefficients were ranked by coefficient magnitude, and the top 15 genes from each model were retained. Models demonstrating a significant association with PFS in the discovery cohort (multivariable Cox PH, p < 0.05) were retained, and genes that appeared among the top 15 predictors across 6 of the 11 models were prioritized, yielding a final outcome-associated spatial DEG signature comprising 15 genes. For each patient, a composite signature score was calculated as the linear predictor derived from the elastic net model, defined as the weighted sum of scaled pseudobulk expression values for the 15 selected genes using their corresponding regression coefficients. Signature scores were standardized using a z-score transformation based on the mean and standard deviation of the discovery cohort. The prognostic performance of the resulting signature was first evaluated in the discovery cohort and subsequently assessed in the independent validation cohort (N = 71). Associations between the signature score and PFS were tested using multivariable Cox proportional hazards models adjusted for age, BMI, receipt of frontline ASCT, and site-batch. For visualization, patients were stratified into high- and low-risk groups using an optimal cut point value based on the standardized signature score in the discovery cohort, and Kaplan-Meier curves were generated to compare survival outcomes between groups.

### Extramedullary myeloma biopsy sample collection and sequencing

Spatial transcriptomic profiling was performed using 10X Genomics V-HD Spatial Gene Expression for FFPE platform according to manufacturers’ instructions. FFPE tissue blocks were sectioned with 5 µm thickness onto negatively charged microscope glass slides (ThermoFisher 12-550-15). The sections were deparaffinized and decrosslinked. Human whole transcriptome probe panels consisting of specific probes for each targeted gene were then added to the tissue. These probe pairs hybridize to their gene target and ligate to one another. Tissue slides and Visium CytAssist Spatial Gene Expression Slides were loaded into the Visium CytAssist instrument, where they were brought into proximity with one another. Each Visium CytAssist HD Slide contains about 11 million Capture Areas (2×2 um) with barcoded spots that include oligonucleotides required to capture gene expression probes. Gene expression probes were released from the tissue upon CytAssist Enabled RNA Digestion & Tissue Removal, enabling capture by the spatially barcoded oligonucleotides present on the Visium slide surface. All the probes captured on a specific spot share a common Spatial Barcode. Libraries were subsequently generated from the probes and sequenced, and the Spatial Barcodes were used to associate the reads back to the tissue section images for spatial mapping of gene expression. Libraries were sequenced on a platform. Sequencing was performed on an Illumina NovaSeq sequencer. *Extramedullary myeloma Visium HD, processing, and quality control* HD sequencing data for extramedullary MM were processed and underwent QC using the same procedures as the bone marrow samples. Sequencing data were processed using Space Ranger (v3.0.1, 10x Genomics Inc.) to demultiplex sequencing reads into FASTQ files, align reads to the human reference genome (GRCh38 2020-A), and generate gene-by-voxel UMI count matrices. For consistency with the bone marrow analyses, we used the 16 µm x 16 µm voxels to maximize the number of QC-passing voxel size. Voxels were retained if passing QC thresholds of at least 100 UMIs and less than 20% of UMIs mapping to mitochondrial genes.

### Summary of extramedullary myeloma Visium HD analyses

Extramedullary V-HD data underwent the same sequence of analyses as described for the bone marrow. Voxels were deconvolved using the same analytical pipeline and reference dataset as was used for bone marrow. Enrichment for cell type gene scores was assessed using the same approach. Cellular neighborhoods were annotated using the same KNN-based methodology. Differential expression was executed using the MAST approach. Spatial concordance between non-canonical Wnt ligand expression and downstream pathway expression was assessed using the same methodology. For T cells, the pro-cytotoxicity ligands were *CCL17, CXCL10, CXCL9, CCL19, CCL5,* and *CCL8*, and the genes used to indicate T cell cytotoxicity were *CD3D, CD3E, CD3G, CD8A, CD8B, NKG7, GNLY, PRF1, GZMB, GZMH, GZMK, CTSW, KLRD1, KLRK1, KLRC1, KLRC2, IFNG, TBX21, EOMES,* and *CX3CR1*.

## Acknowledgments

The authors would like to acknowledge funding support from the Myeloma Solutions Fund (MB, DJO, MEM, WCP), National Cancer Institute (NCI) R01 CA252222, R01 CA244899, R01 CA262754, K12 CA270375, CA196521 (SP), and the Multiple Myeloma Research Foundation (MB, SP).

