## Supplemental Figures and Legends for "Spatial multi-omics of multiple myeloma uncovers niche-dependent pro-myeloma and immunosuppressive signaling in the bone marrow and extramedullary lesions"

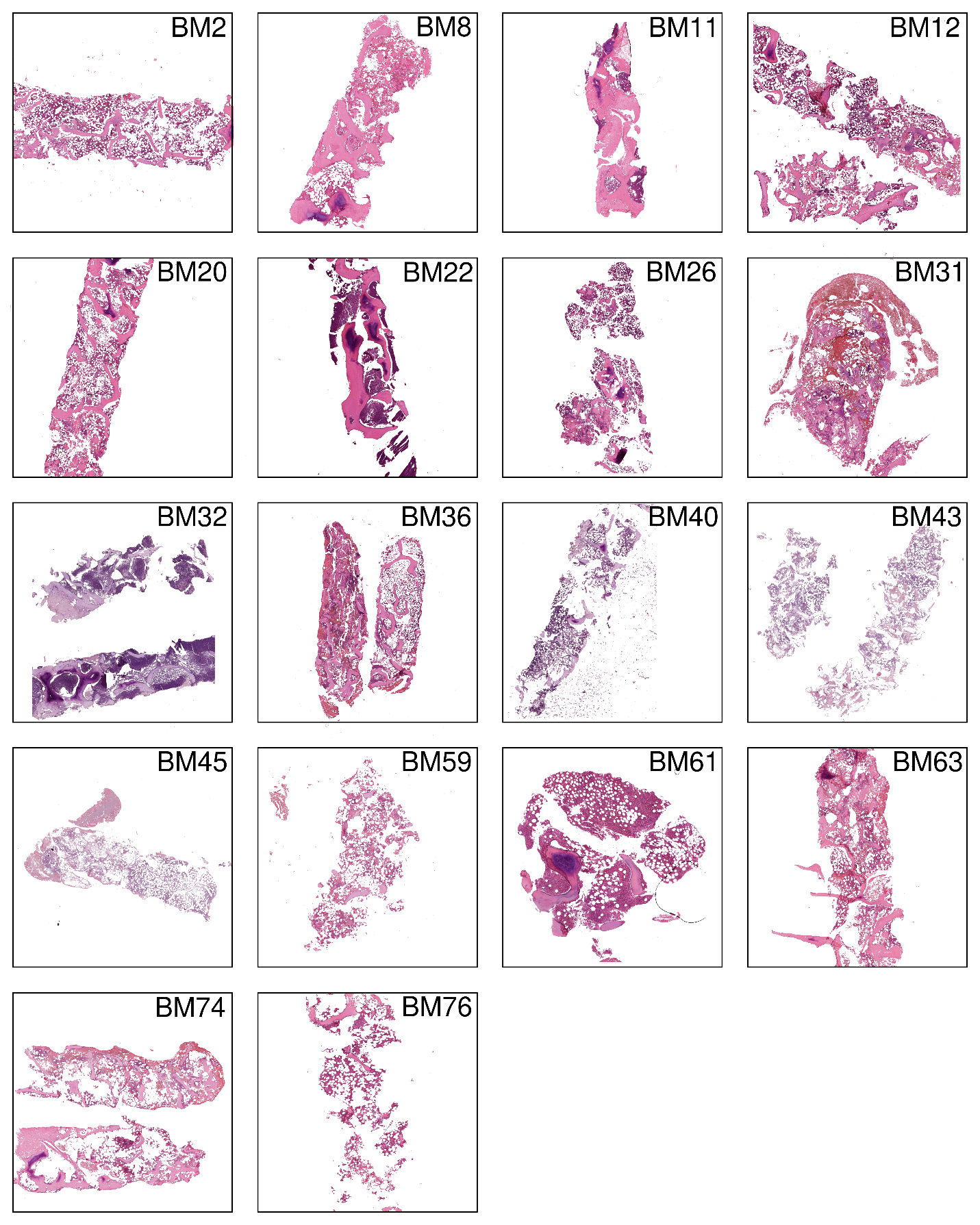

**Supplemental Figure 1. Hematoxylin and eosin staining of bone marrow biopsies in the Visium HD cohort.** Hematoxylin and eosin staining of the bone marrow biopsy for the Visium HD cohort. BM0 and BM28 did not have staining images available, BM17 is shown in Figure 2.

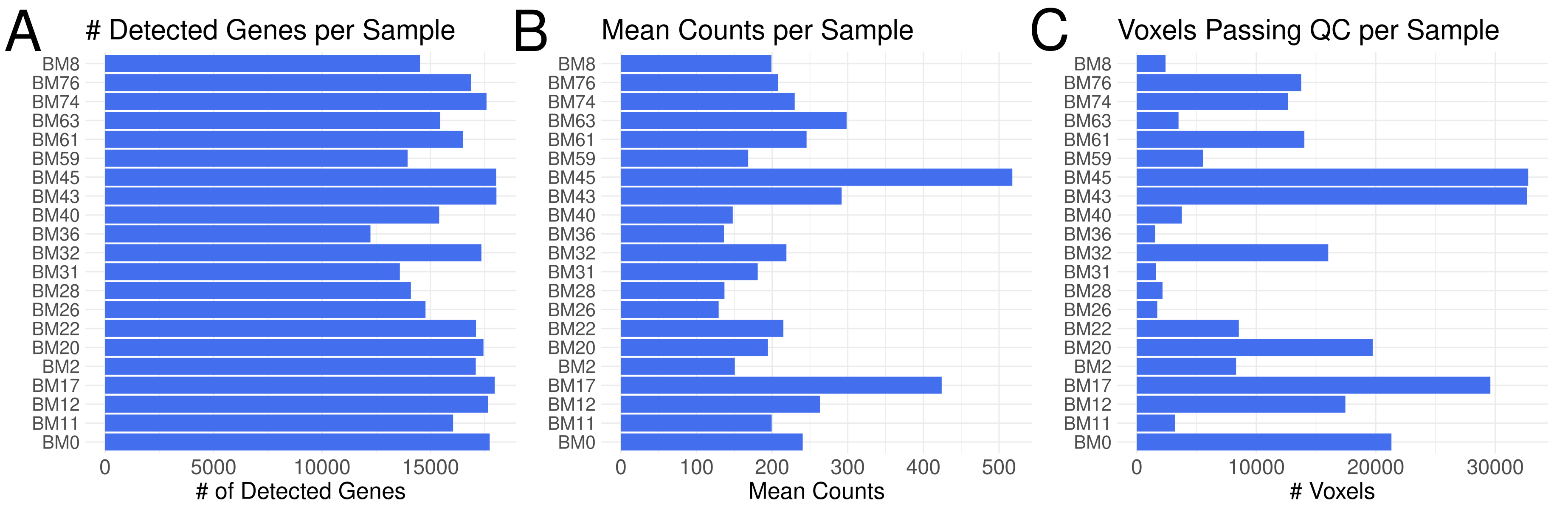

**Supplemental Figure 2. Quality metrics for the Visium HD cohort. (A-C)** Bar plots displaying the total number of unique genes detected per sample (A), the mean counts per voxel for each sample (B), and the number of voxels per sample that passed quality filtering of ≥ 100 counts and ≤ 20% mtRNA. QC-quality control.

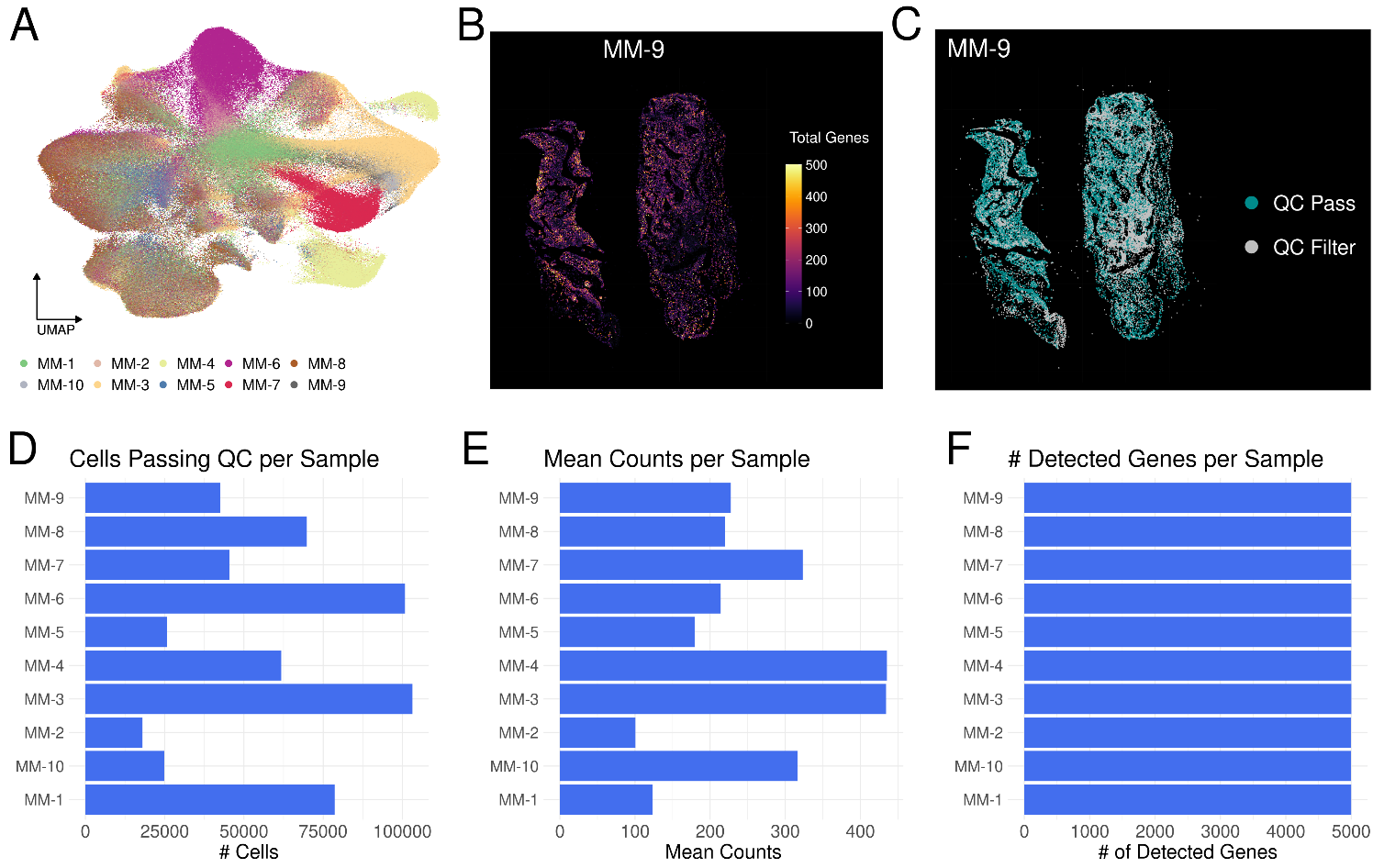

**Supplemental Figure 3. Quality metrics for the Xenium cohort. (A)** Uniform manifold approximation and projection plot displaying the cells from 10 MM bone marrow biopsies published by Yip et al. Points are colored by patient. **(C)** Representative spatial dot plot displaying the total number of unique gene detected per cell for sample MM-9. **(D)** Representative spatial dot plot for cells passing quality control (teal) or filtered out from downstream analyses (gray). Consistent with Yip *et al*., quality controls filters were ≥ 60 total counts per cell and ≥ 50 unique genes per cell. **(D-F)** Bar plots displaying the number of cells per sample that passed quality filtering (D), the mean counts per voxel for each sample (E), and the total number of unique genes detected per sample (F). QC-quality control.

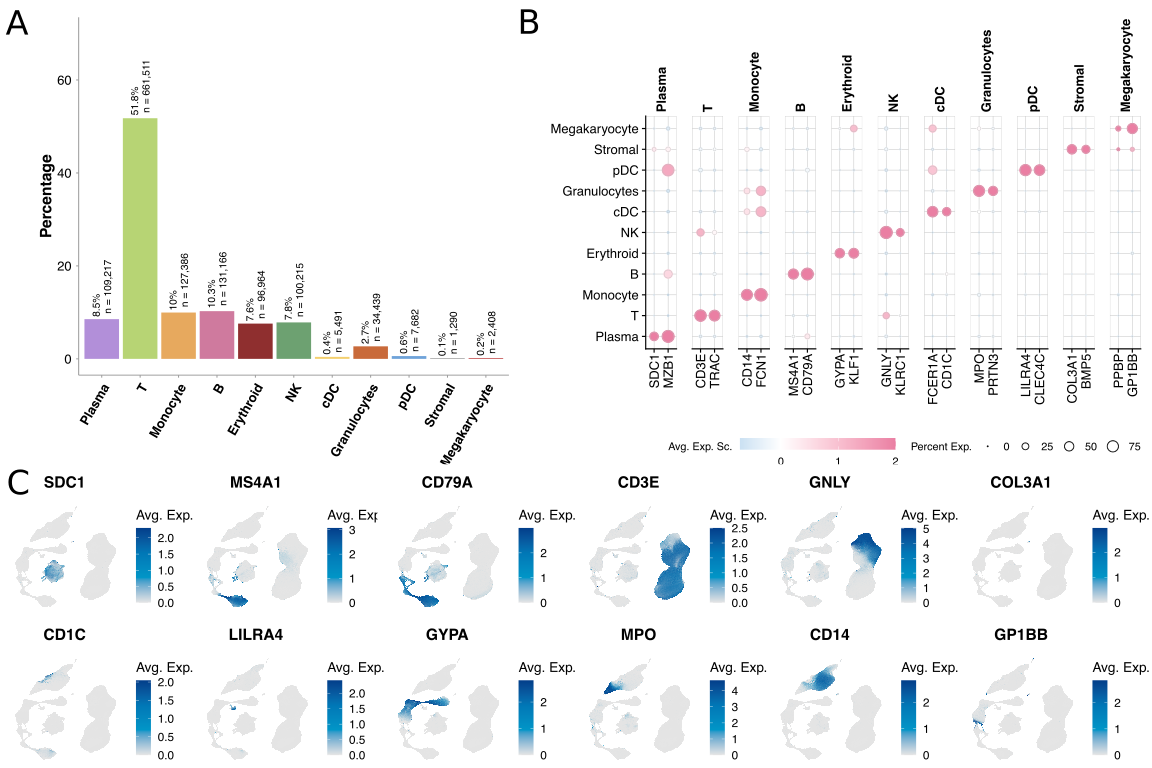

**Supplemental Figure 4. Annotation of cells in the single-cell RNA sequencing cohort. (A)** Bar plots displaying the percentage of cells by cell type in the Pilcher et al. single cell RNA sequencing dataset. **(B)** Dot plot displaying the average normalized and scaled lineage marker gene expression by cell type. **(C)** Uniform manifold approximation and projection plots displaying the normalized expression of canonical lineage genes used to annotate cell types. Avg. Exp. Sc.-Average expression score,

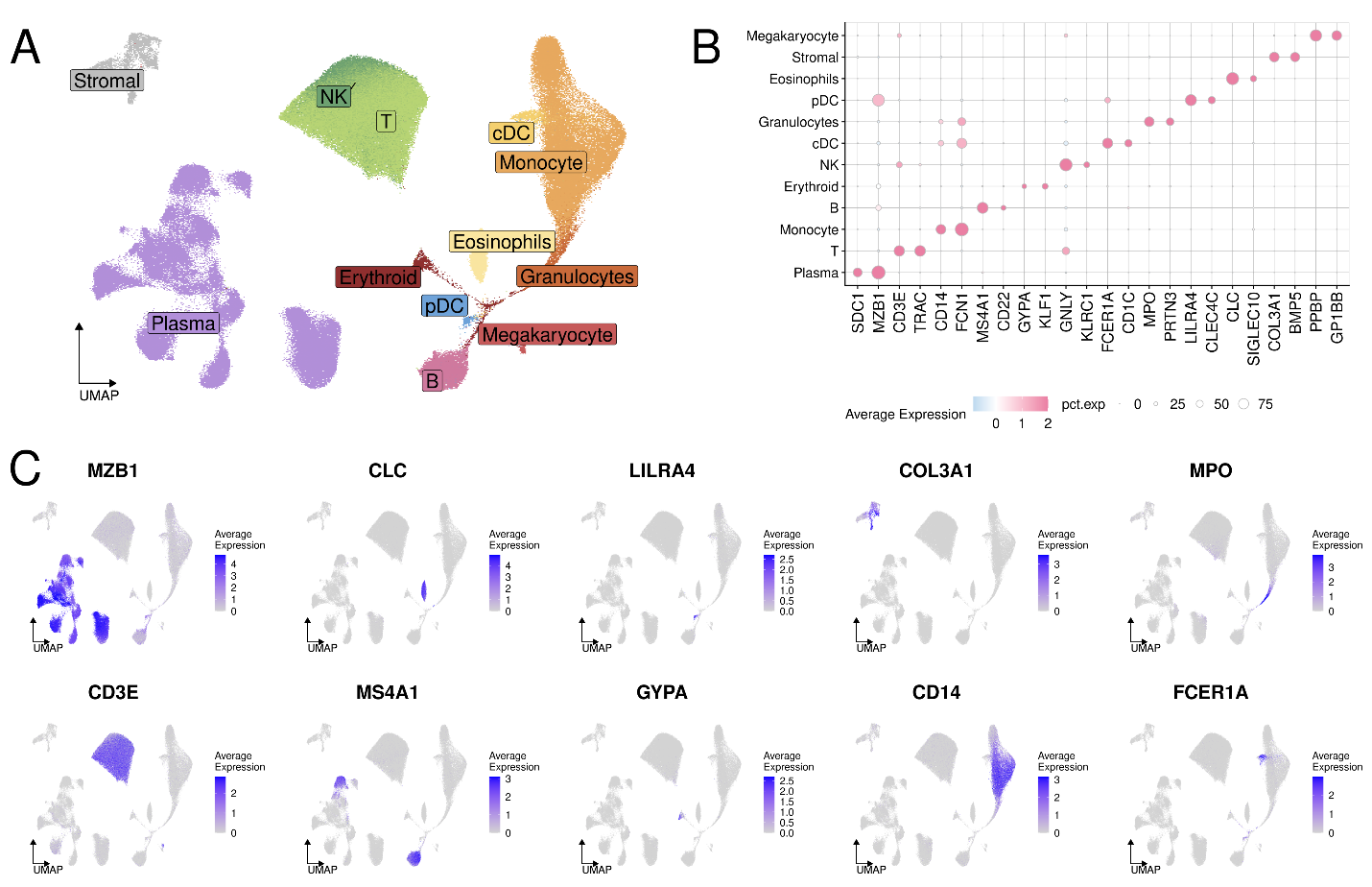

**Supplemental Figure 5. Annotations of cells in the single-cell RNA sequencing deconvolution reference dataset.** To deconvolve the cellular composition of voxels in the Visium HD cohort we generated a custom reference single-cell RNA sequencing dataset. The reference is primarily composed of unsorted bone marrow biopsies from GSE. During quality control we noted that stromal cells and eosinophils were well-detected in the Visium HD data; cell types that are typically not well captured in single-cell RNA sequencing data. Therefore, we merged sorted stromal cells from GSE, and eosinophils from GSE to appropriately deconvolve the Visium HD data. **(A)** Uniform manifold approximation and projection plot with points colored by cell types in the merged reference dataset. **(B)** Dot plot displaying the average normalized and scaled lineage marker gene expression by cell type. **(C)** Uniform manifold approximation and projection plots displaying the normalized expression of canonical lineage genes used to annotate cell types.

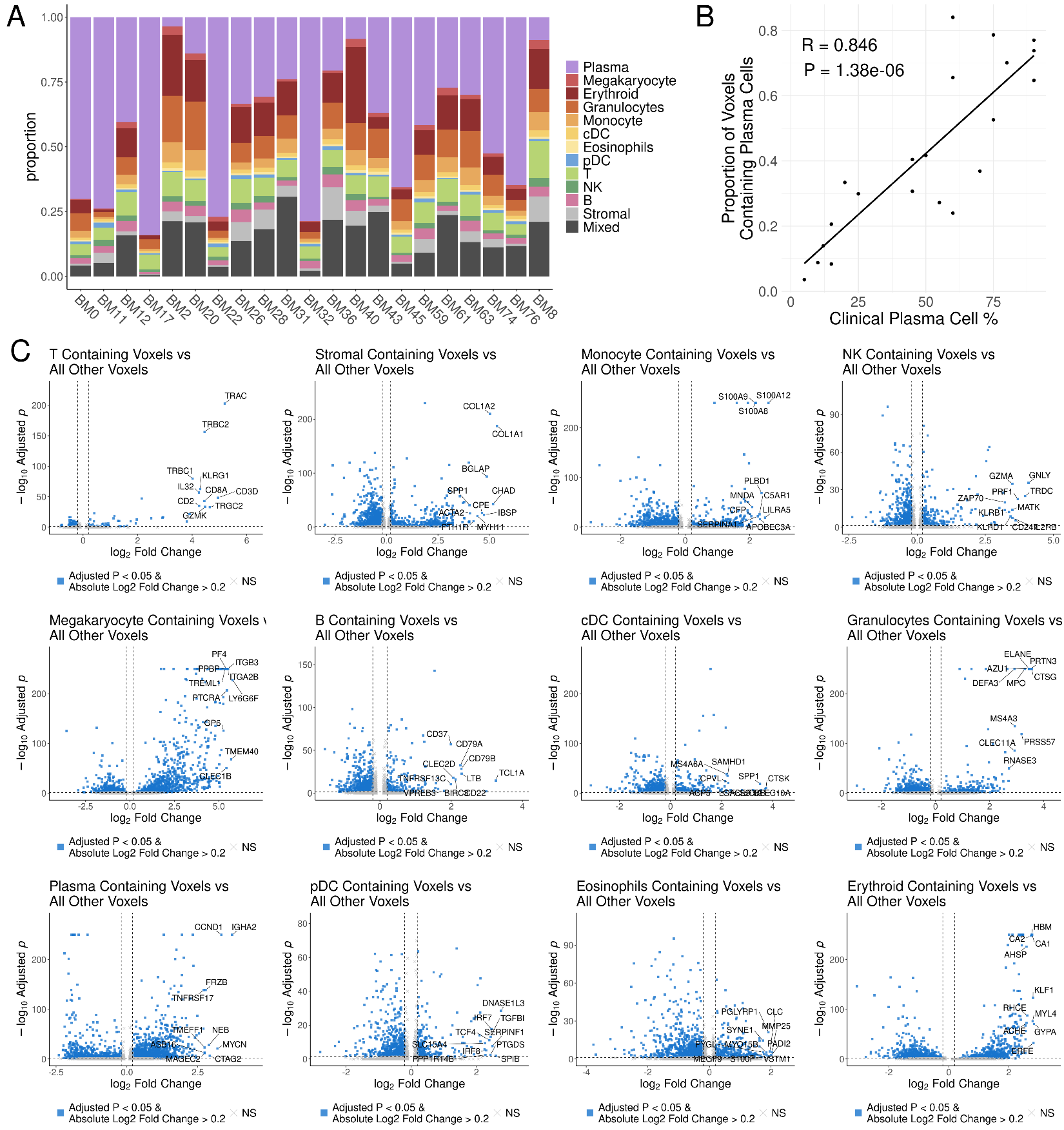

**Supplemental Figure 6. Voxel deconvolution in Visium HD data accurately identifies contributing cell types. (A)** Bar plot displaying the proportion of cell type annotations for each sample in the Visium HD cohort. Voxels that did not meet deconvolution threshold criteria for any cell type were labeled “Mixed”. **(B)** Dot plot displaying the estimated plasma cell percentage by clinical pathological examination (X axis) versus the proportion of voxels annotated as plasma cell-containing (Y axis). R and P value calculated by Pearson correlation test. **(C)** Volcano plots for each cell type testing for enrichment for canonical lineage genes. Differential expression was tested using a hurdle model as implemented by the MAST method in Seurat’s FindMakers function with a significance cutoff of multiple comparisons adjusted p value < 0.05.

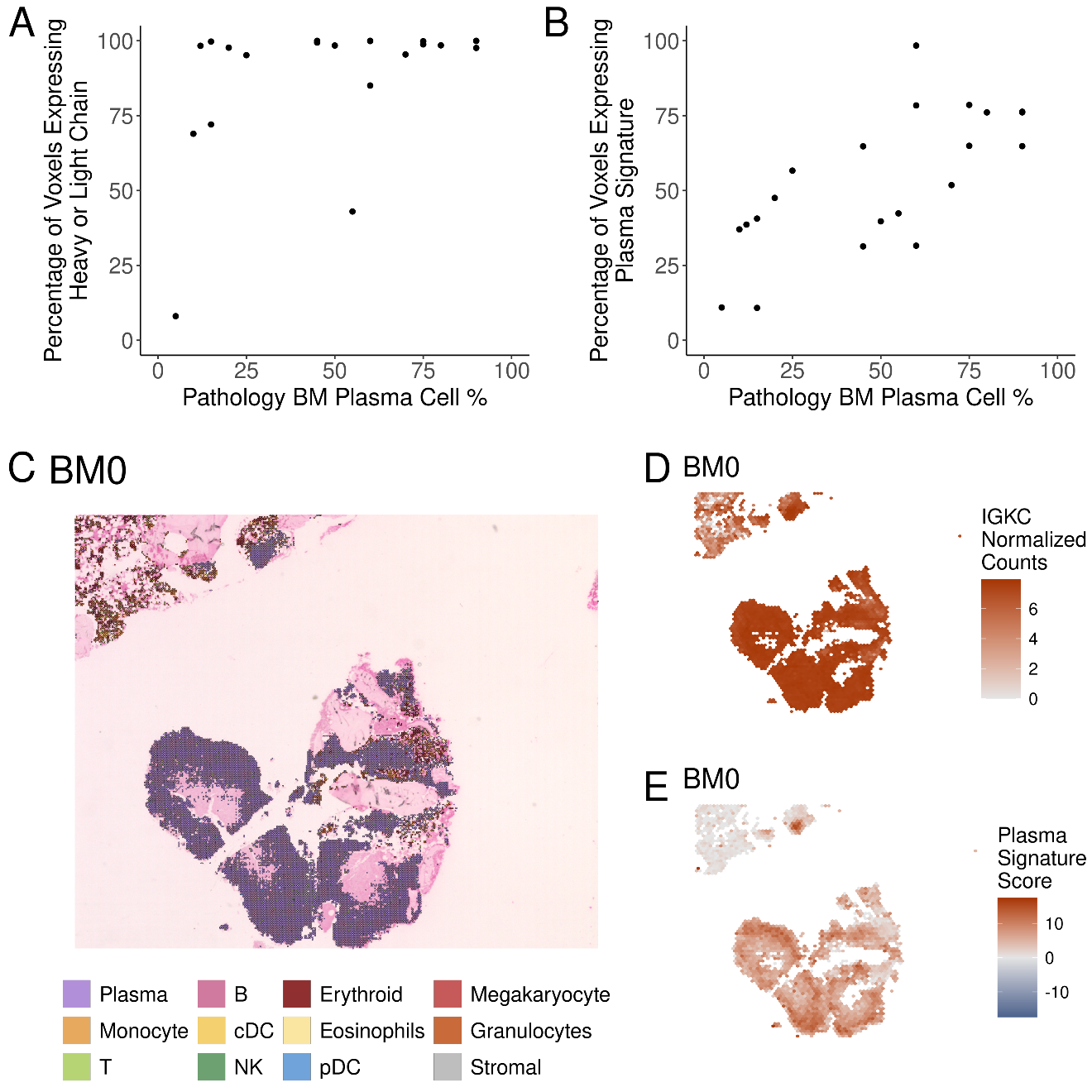

**Supplemental Figure 7. Voxel deconvolution mitigates broad detection of individual genes. (A)** Dot plot displaying the percentage of voxels with detection of a patients’ involved heavy or light immunoglobulin chain (Y axis) versus the clinical pathology estimate of bone marrow plasma cell percentage (X axis). Findings are consistent with prior studies, such as Gong et al., that find highly abundant genes like the immunoglobulin genes to be over-represented in sequencing-based spatial transcriptomics, even in samples from healthy controls where plasma cells are relatively rare. **(B)** Dot plot displaying the percentage of voxels with detection of the plasma cell gene signature used in this study (Y axis) versus the clinical pathology estimate of bone marrow plasma cell percentage (X axis). See methods for the plasma cell gene list. **(C)** Representative spatial scatterpie plot showing the annotation of voxels in BM0. Voxels are colored by cell type, when multiple cell types are present in a given voxel the scatterpie plot is equally split and colored by the cell types present. **(D-E)** Representative spatial hexbin plots displaying the density of *IGKC* in BM0 which is a kappa light chain myeloma (D), versus the density of the plasma cell expression signature as in B (E).

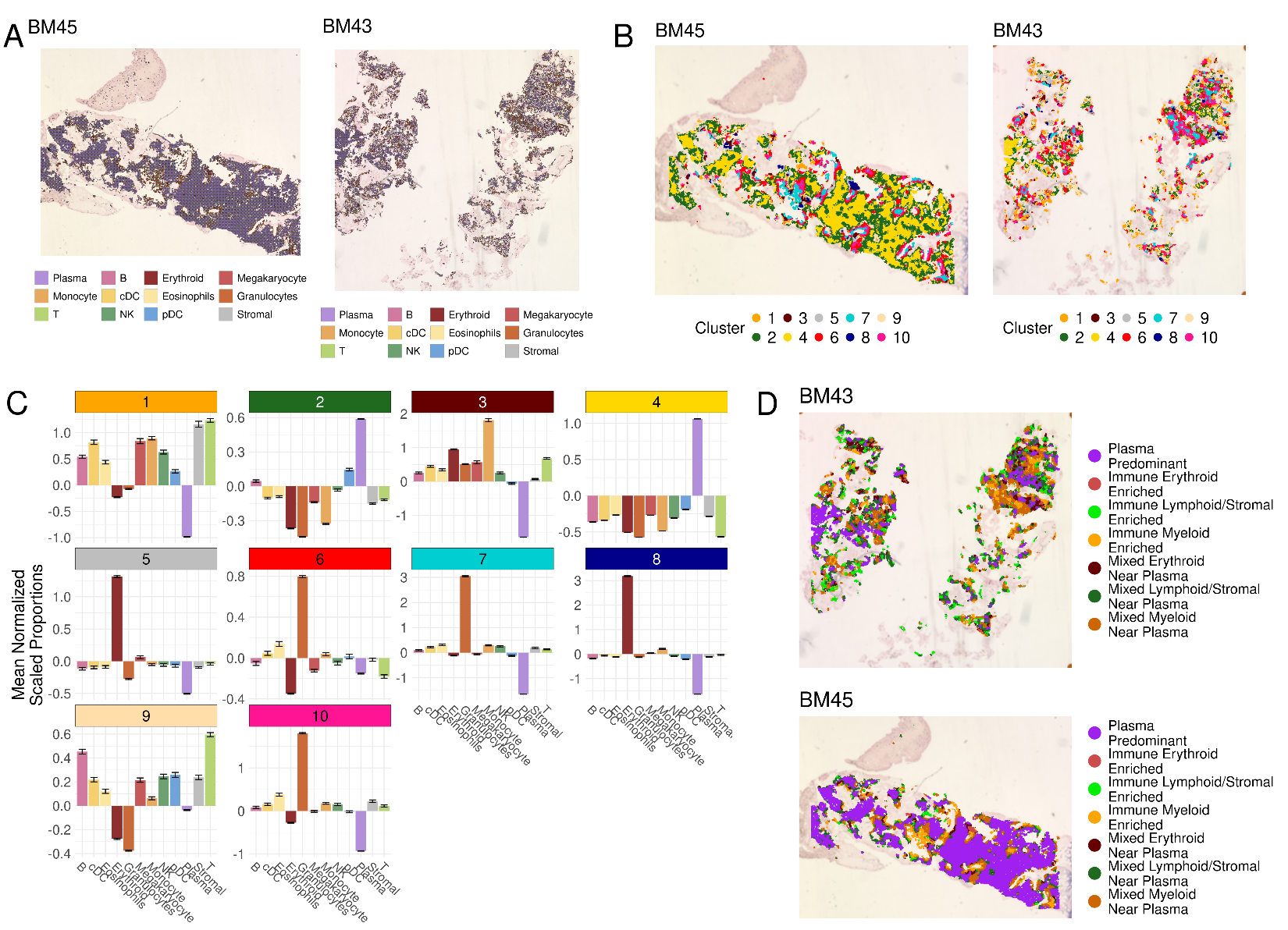

**Supplemental Figure 8. Neighborhood annotation in the Visium HD cohort. (A)** Representative spatial scatterpie plots showing the annotation of voxels in a sample with relatively high plasma cell burden (BM45, left) and relatively low plasma cell burden (BM43, right). Voxels are colored by cell type, when multiple cell types are present in a given voxel the scatterpie plot is equally split and colored by the cell types present. **(B)** Representative spatial dot plot displaying the unsupervised K-nearest neighbor (KNN) clusters. For each voxel, the proportion of cell types within a 50 µm radius was quantified and used for KNN clustering of voxels by proximal cell composition. **(C)** Bar plots depicting the relative abundance of cell types in the ten KNN clusters. **(D)** Representative spatial dot plot displaying the neighborhood annotations. KNN clusters were annotated based on the enriched cell types as displayed in C.

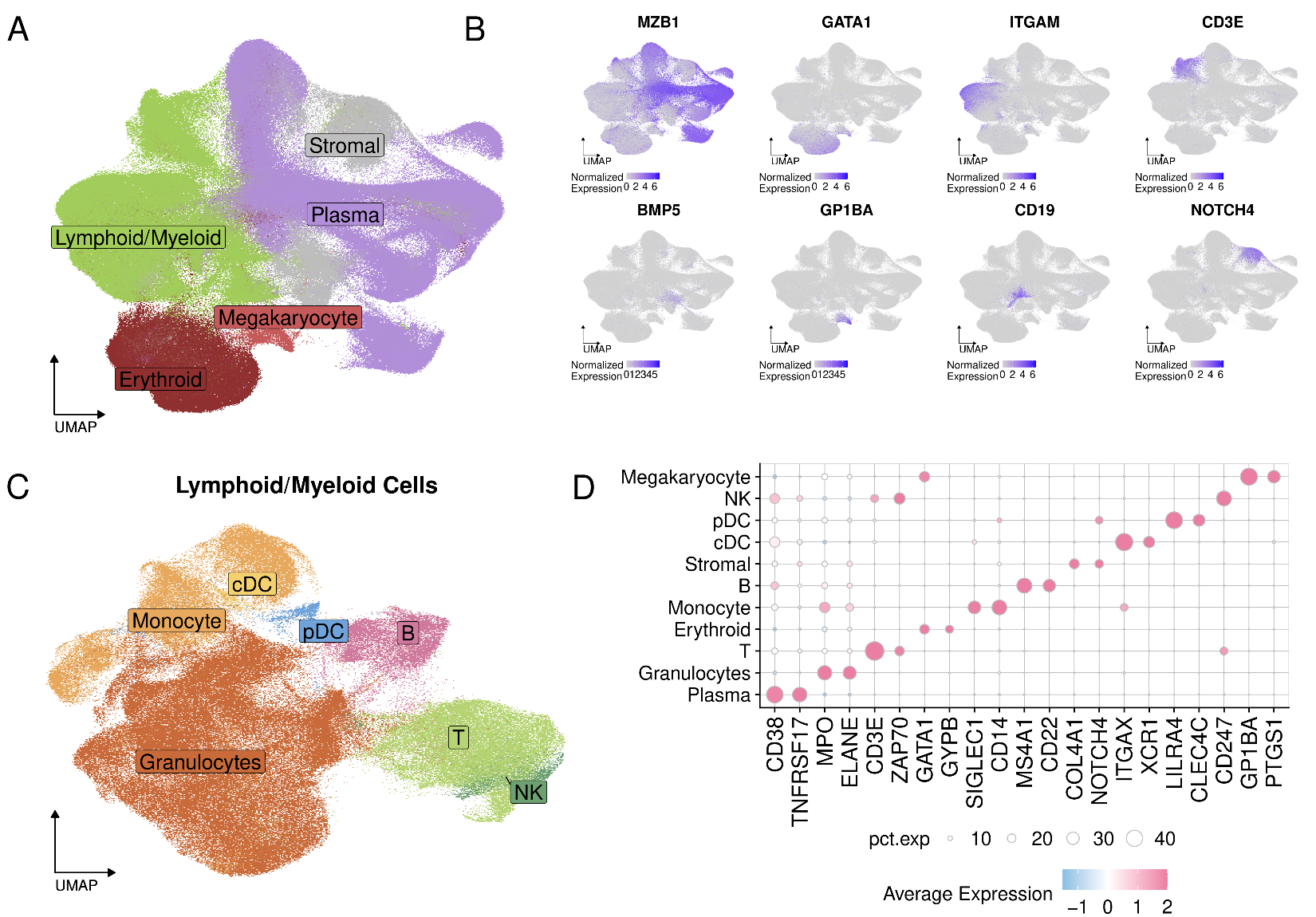

**Supplemental Figure 9. Annotations of cells in the Xenium cohort.** **(A)** Uniform manifold approximation and projection plot with points colored by broad cell types annotated in the Xenium data. **(B)** Uniform manifold approximation and projection plots displaying the normalized expression of canonical lineage genes used to annotate broad cell types. **(C)** Uniform manifold approximation and projection plot of lymphoid and myeloid cells with points colored by cell type annotation harmonized to the cell types annotated in Visium data. Lymphoid and myeloid cells did not form discrete clusters at the dataset level; Lymphoid and myeloid cells were independently subclustered and annotated using canonical lineage markers. **(D)** Dot plot displaying the average normalized and scaled lineage marker gene expression by cell type.

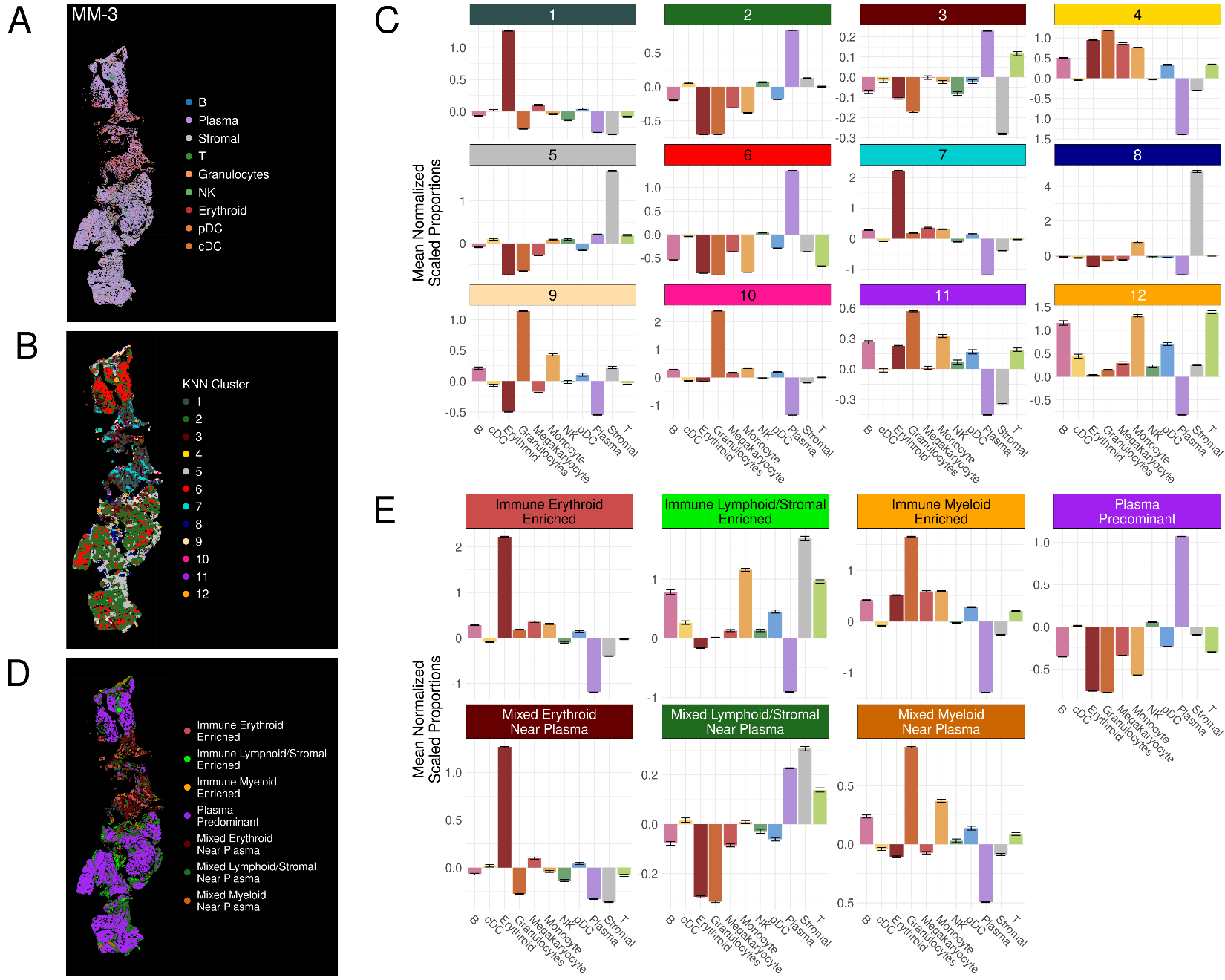

**Supplemental Figure 10. Neighborhood annotation in the Xenium cohort.** Cells in the Xenium cohort underwent the same K-nearest neighbor (KNN) clustering analysis as the Visium cohort. **(A)** Representative spatial dot plot showing the cell type annotations in MM-3. **(B)** Representative spatial dot plot displaying the unsupervised KNN clusters. For each voxel, the proportion of cell types within a 50 µm radius was quantified and used for KNN clustering of voxels by proximal cell composition. **(C)** Bar plots depicting the relative abundance of cell types in the twelve KNN clusters. **(D)** Representative spatial dot plot displaying the neighborhood annotations. KNN clusters were annotated based on the enriched cell types as displayed in C. **(E)** Bar plots depicting the relative abundance of cell types in the seven annotated neighborhoods.

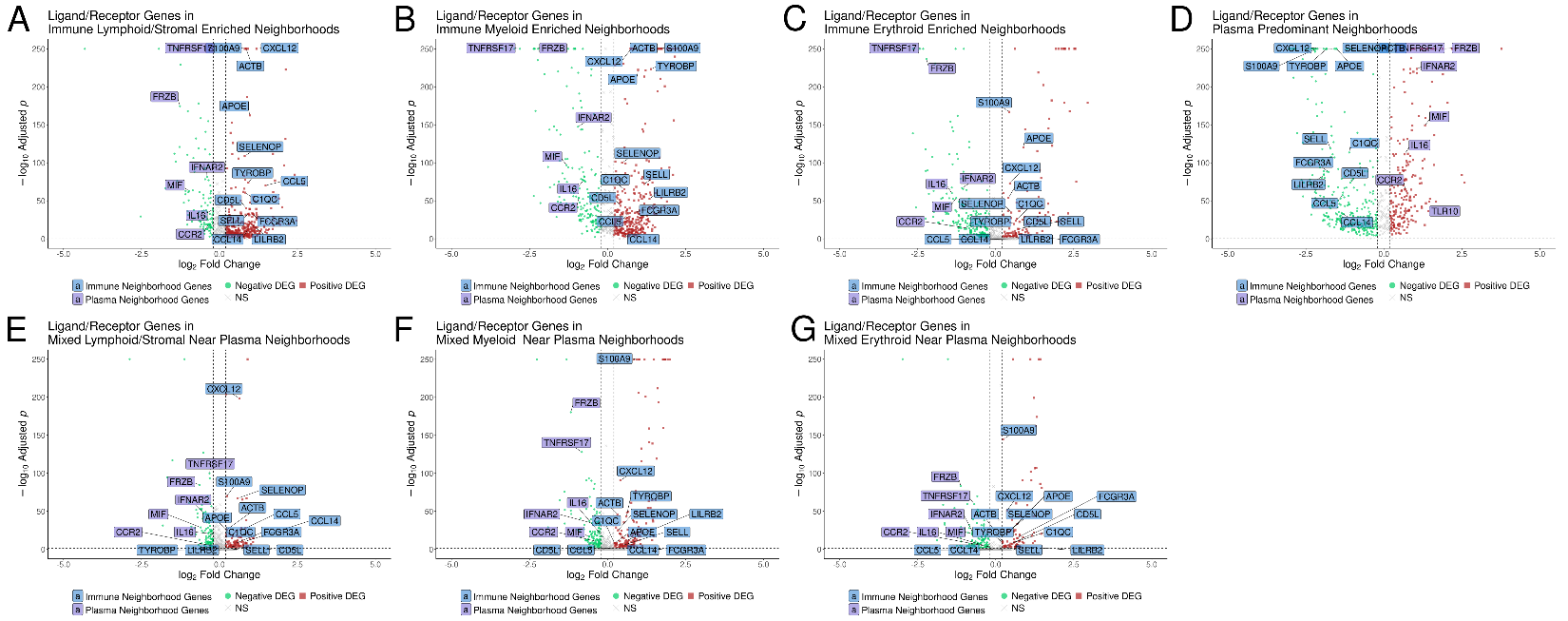

**Supplemental Figure 11. Non-malignant neighborhoods display similar expression of ligand and receptor genes. (A-G)** Volcano plots for each neighborhood in Visium HD data testing for enrichment for ligand and receptor genes. Differential expression was tested using a hurdle model as implemented by the MAST method in Seurat’s FindMakers function with a significance cutoff of multiple comparisons adjusted p value < 0.05. Gene labels are colored by whether they were consistently observed to be enriched in the six types of non-malignant neighborhoods or in the plasma-predominant neighborhoods.

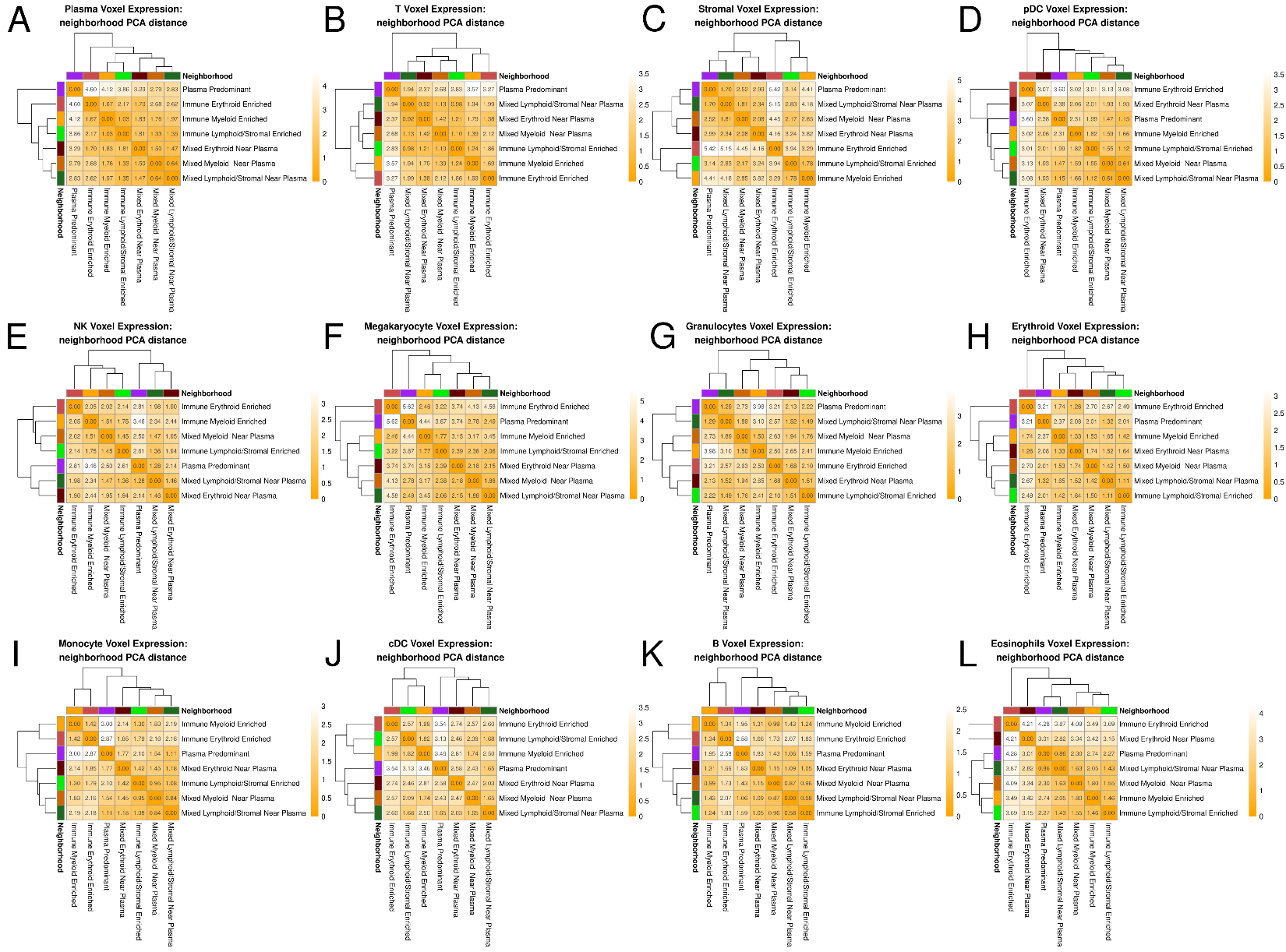

**Supplemental Figure 12. Cells within non-malignant neighborhoods have similar transcriptomic profiles in the Visium HD cohort. (A-L)** Heatmaps displaying the average distance in principal component analysis space for each cell type across neighborhoods. For each cell type, principal component analysis was performed on the top 200 highly variable cell type genes. Then the Euclidean distance was calculated on the first 50 principal components to provide a metric of transcriptome similarity across neighborhoods where lower distance indicates more similar expression.

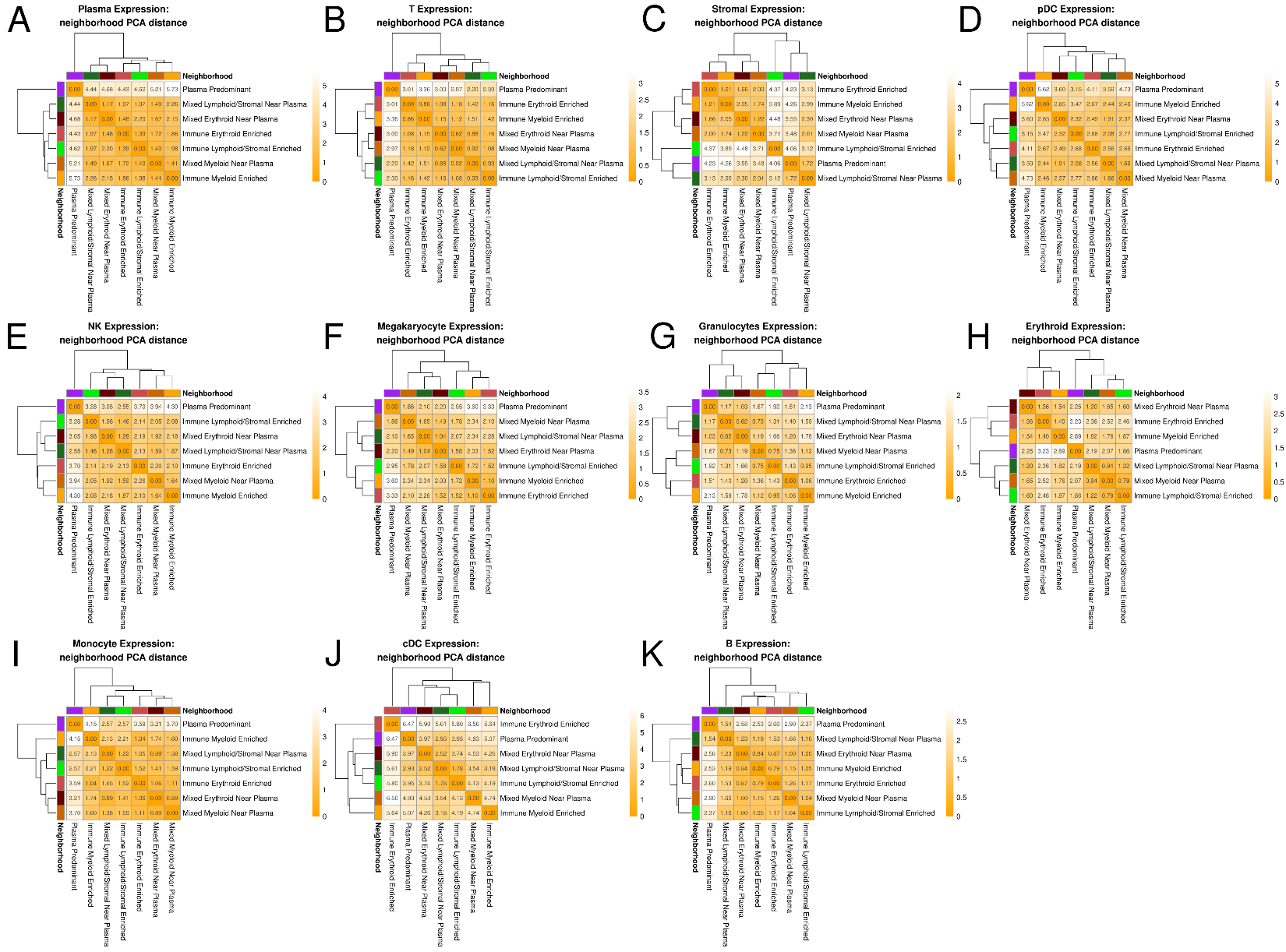

**Supplemental Figure 13. Cells within non-malignant neighborhoods have similar transcriptomic profiles in the Xenium cohort. (A-K)** Heatmaps displaying the average distance in principal component analysis space for each cell type across neighborhoods. For each cell type, principal component analysis was performed on the top 200 highly variable cell type genes. Then the Euclidean distance was calculated on the first 50 principal components to provide a metric of transcriptome similarity across neighborhoods where lower distance indicates more similar expression.

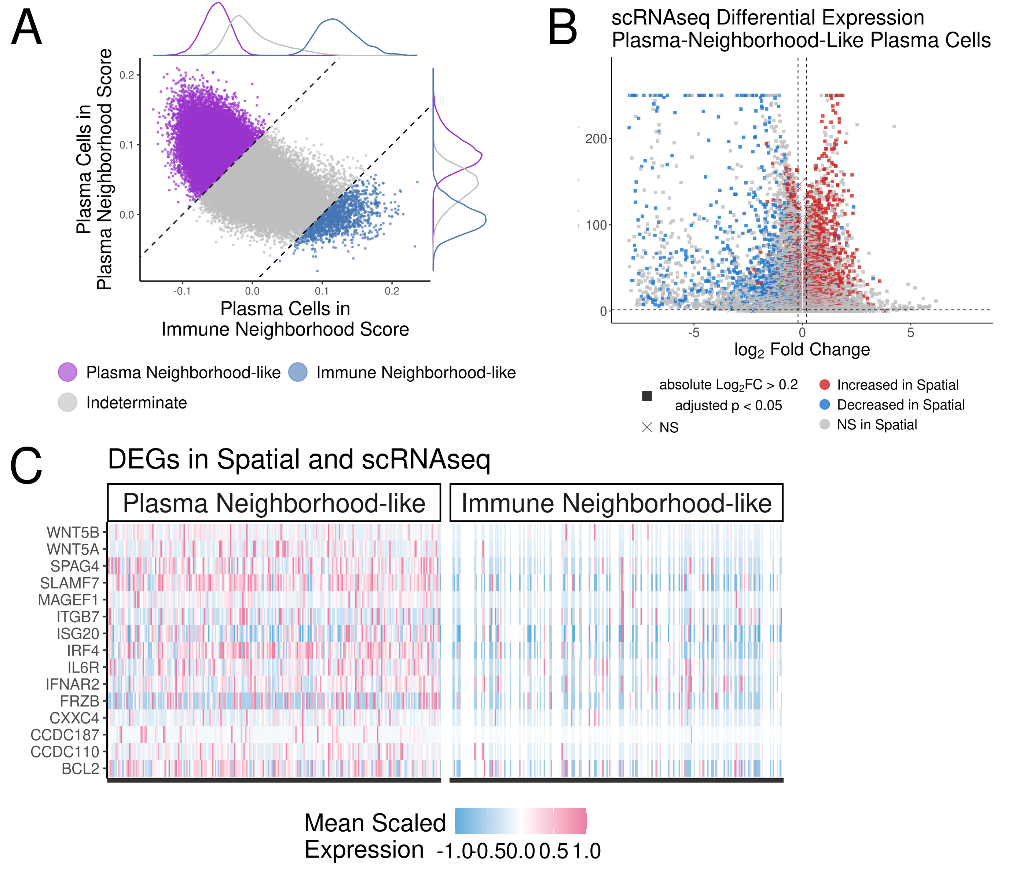

**Supplemental Figure 14. Identification of plasma cells in single-cell RNA sequencing with neighborhood-like gene expression profiles. (A)** Dot plot depicting the annotation of plasma neighborhood-like and immune neighborhood-like plasma cells in the single-cell RNA sequencing cohort. The spatially differentially expressed genes of plasma cells were used to calculate plasma and immune neighborhood-like scores using the Seurat AddModuleScore function. A differentiation threshold of 0.1 was used, with cells below this threshold being annotated as “Indeterminate”. **(B)** Volcano plot comparing plasma neighborhood-like plasma cells to immune neighborhood-like plasma cells. Differential expression was tested using a hurdle model as implemented by the MAST method in Seurat’s FindMakers function with a significance cutoff of multiple comparisons adjusted p value < 0.05. Points are colored by whether the each gene was positively differentially expressed (red), negatively differentially expression (blue) or not differentially expressed in spatial data to display the coherence in gene directionality as well as the enhanced gene detection in single-cell RNA sequencing relative to spatial assays. **(C)** Heatmap displaying the mean normalized scaled expression of genes that were differentially expressed between plasma-neighborhood-like plasma cells versus immune-neighborhood-like plasma cells in spatial analyses.

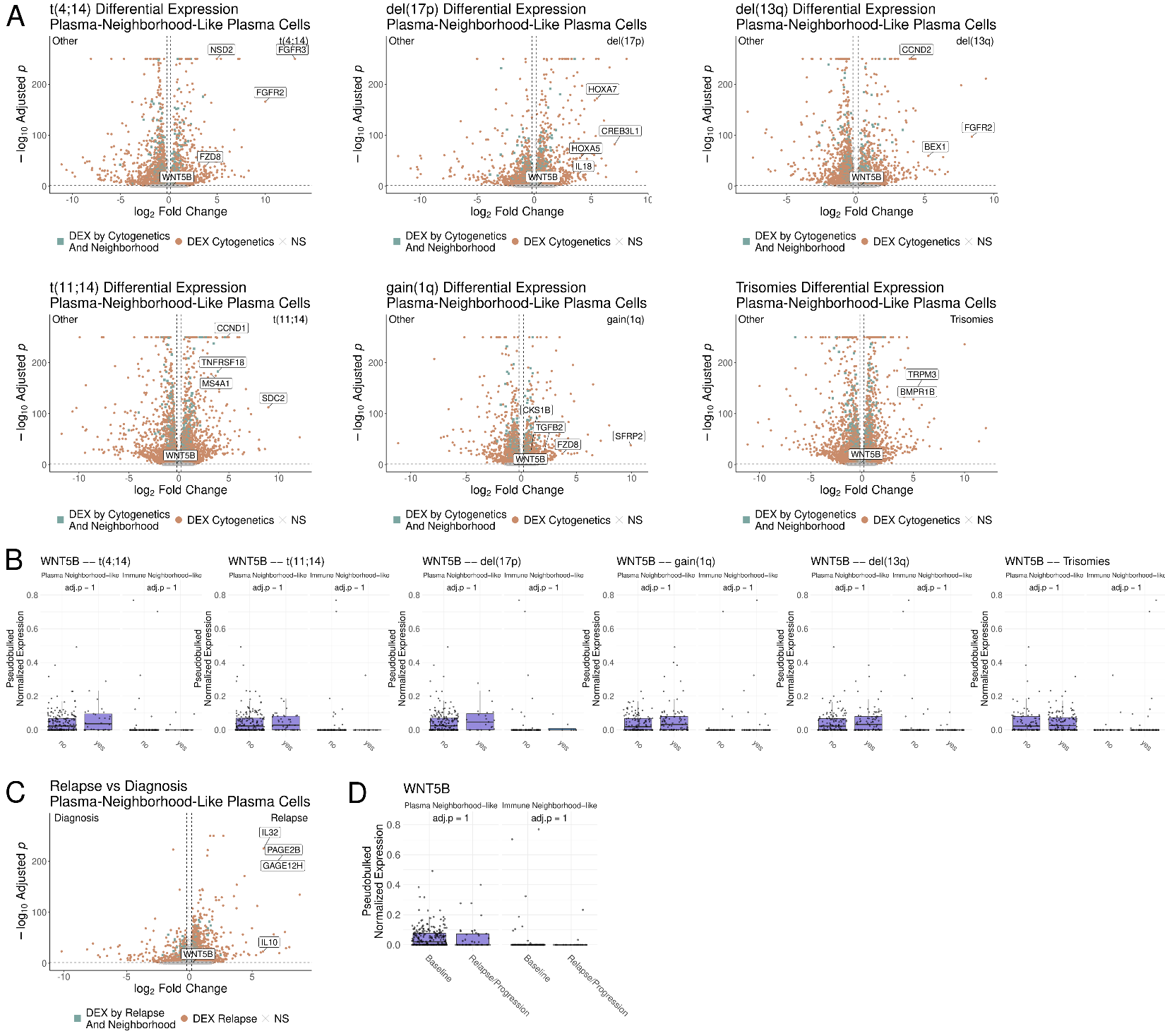

**Supplemental Figure 15. Comparison of plasma neighborhood-like plasma cells across cytogenetics and disease progression finds persistent expression of non-canonical Wnt promoting ligand, WNT5A. (A)** Volcano plot comparing plasma neighborhood-like plasma cells between samples with and without t(4;14), t(11;14), del(17p), gain(1q), del(13q) and odd numbered trisomies. Differential expression was tested using a hurdle model as implemented by the MAST method in Seurat’s FindMakers function with a significance cutoff of multiple comparisons adjusted p value < 0.05. Points are colored in green for genes differentially expressed by cytogenetics and by neighborhood (plasma neighborhood-like versus immune neighborhood-like as in **Fig 3C**), orange for cytogenetics only, and gray for not differentially expressed. **(B)** Box plots displaying the patient-wise pseudobulked normalized expression of *WNT5B* across cytogenetic groups and neighborhood-like groups. **(C)** Volcano plot comparing plasma neighborhood-like plasma cells between samples at relapse versus diagnosis. Differential expression performed as in (A). Points are colored in green for genes differentially expressed by relapse and by neighborhood (plasma neighborhood-like versus immune neighborhood-like as in **Fig 3C**), orange for relapse only, and gray for not differentially expressed. **(B)** Box plots displaying the patient-wise pseudobulked normalized expression of *WNT5B* across diagnosis and relapse and neighborhood-like groups.

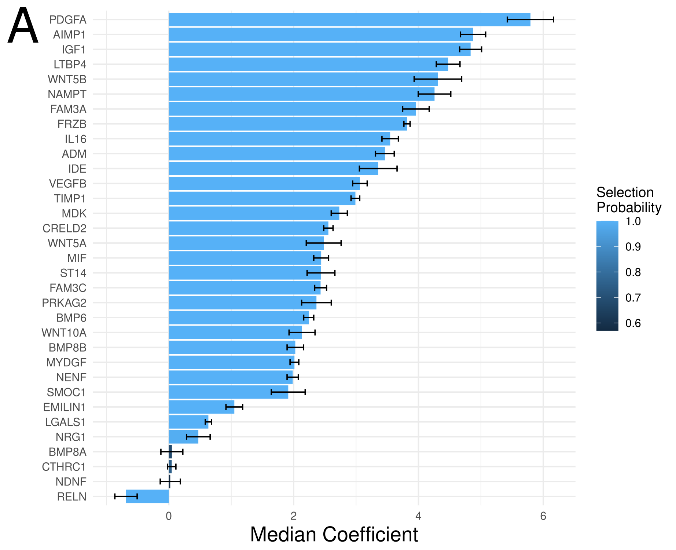

**Supplemental Figure 16. Elastic net regression identifies ligands associated with spatial alteration in T cell expression. (A)** Bar plot showing the ligand genes ranked by their spatial association with altered T cell expression. For each T cell, the total detection of ligand gene expression was calculated within a 50 µm radius. Additionally, a spatial T expression score was calculated from the spatially differentially expressed genes of T cells in plasma-predominant neighborhoods vs immune-enriched neighborhoods. Then, 100 bootstrapped iterations of elastic net regression was performed with the proximal ligand genes expression as a predictor of the spatial T expression score. Ligand genes were ranked based on a median association coefficient calculated from the coefficient of each ligand in the model multiplied by the frequency with which the ligand was included in the elastic net models.

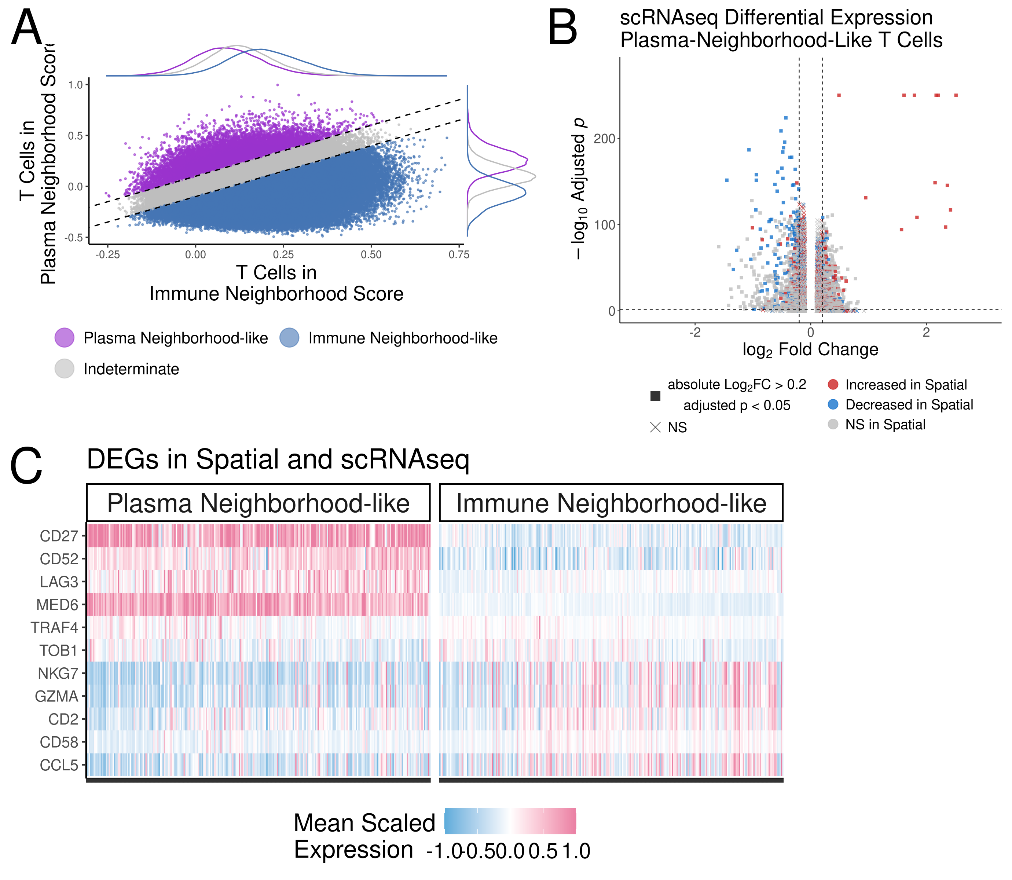

**Supplemental Figure 17. Identification of T cells in single-cell RNA sequencing with neighborhood-like gene expression profiles. (A)** Dot plot depicting the annotation of plasma neighborhood-like and immune neighborhood-like T cells in the single-cell RNA sequencing cohort. The spatially differentially expressed genes of T cells were used to calculate plasma and immune neighborhood-like scores using the Seurat AddModuleScore function. A differentiation threshold of 0.1 was used, with cells below this threshold being annotated as “Indeterminate”. **(B)** Volcano plot comparing plasma neighborhood-like T cells to immune neighborhood-like T cells. Differential expression was tested using a hurdle model as implemented by the MAST method in Seurat’s FindMakers function with a significance cutoff of multiple comparisons adjusted p value < 0.05. Points are colored by whether the each gene was positively differentially expressed (red), negatively differentially expression (blue) or not differentially expressed in spatial data to display the coherence in gene directionality as well as the enhanced gene detection in single-cell RNA sequencing relative to spatial assays. **(C)** Heatmap displaying the mean normalized scaled expression of genes that were differentially expressed between plasma-neighborhood-like T cells versus immune-neighborhood-like T cells in spatial analyses.

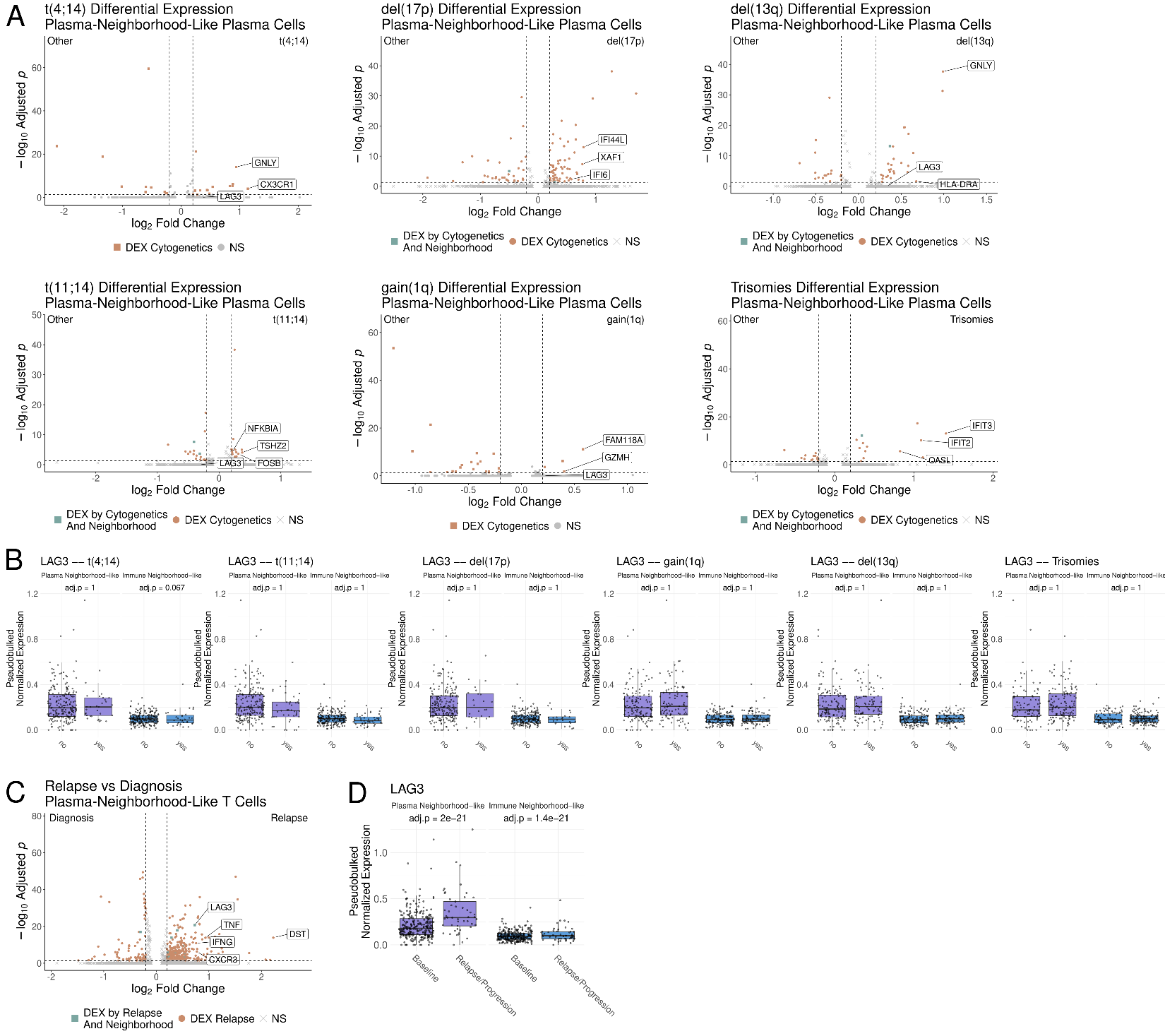

**Supplemental Figure 18. Comparison of plasma neighborhood-like T cells across cytogenetics and disease progression finds persistent expression of LAG3 across cytogenetics and increase expression at relapse. (A)** Volcano plot comparing plasma neighborhood-like T cells between samples with and without t(4;14), t(11;14), del(17p), gain(1q), del(13q) and odd numbered trisomies. Differential expression was tested using a hurdle model as implemented by the MAST method in Seurat’s FindMakers function with a significance cutoff of multiple comparisons adjusted p value < 0.05. Points are colored in green for genes differentially expressed by cytogenetics and by neighborhood (plasma neighborhood-like versus immune neighborhood-like as in **Fig 4D**), orange for cytogenetics only, and gray for not differentially expressed. **(B)** Box plots displaying the patient-wise pseudobulked normalized expression of *LAG3* across cytogenetic groups and neighborhood-like groups. **(C)** Volcano plot comparing plasma neighborhood-like T cells between samples at relapse versus diagnosis. Differential expression performed as in (A). Points are colored in green for genes differentially expressed by relapse and by neighborhood (plasma neighborhood-like versus immune neighborhood-like as in **Fig 4D**), orange for relapse only, and gray for not differentially expressed. **(B)** Box plots displaying the patient-wise pseudobulked normalized expression of *LAG3* across diagnosis and relapse and neighborhood-like groups.

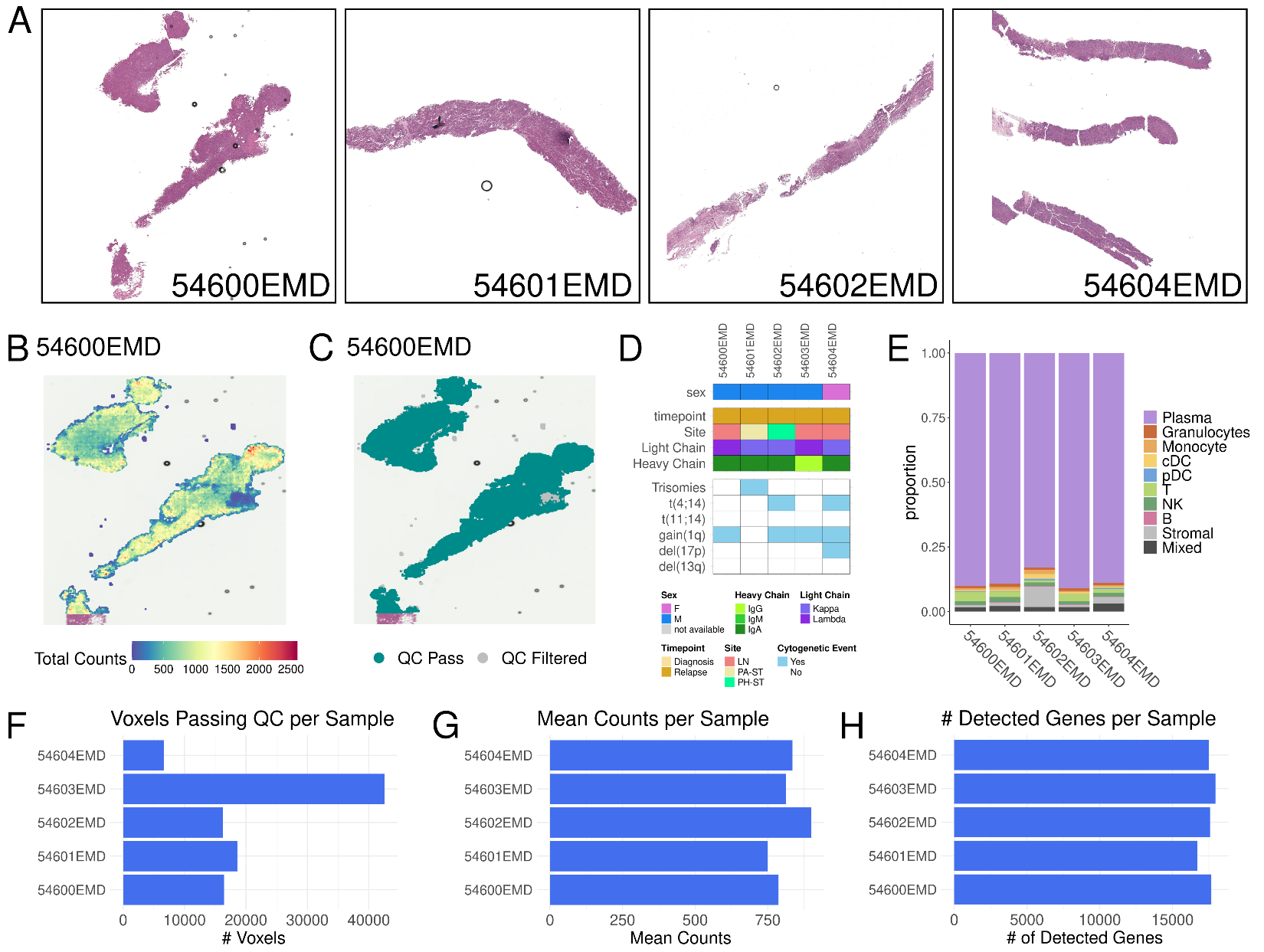

**Supplemental Figure 19. Quality metrics for the extramedullary myeloma Visium HD cohort. (A)** Hematoxylin and eosin staining of the extramedullary (EM) myeloma biopsies for the Visium HD EM cohort. 546003EMD is shown in Figure 6. **(B)** Representative spatial dot plot displaying the total counts detected per cell for sample 54600EMD. **(C)** Representative spatial dot plot for cells passing quality control (teal) or filtered out from downstream analyses (gray). Quality controls filters were ≥ 100 total counts per cell and ≤ 20% mtRNA, consistent with the bone marrow analysis. **(D)** Metadata heatmap showing the clinical and cytogenetic features of the EM samples profiles by Visium-HD. **(E)** Bar plot displaying the proportion of voxel annotations by sample using the same deconvolution approach as the bone marrow analyses. Voxels that did not meet deconvolution thresholds for any cell type were labeled “Mixed”. **(F-H)** Bar plots displaying the number of voxels per sample that passed quality filtering (F), the mean counts per voxel for each sample (G), and the total number of unique genes detected per sample (H). QC-quality control. LN-lymph node, PA-ST-periarticular soft tissue, PH-ST-perihepatic soft tissue.

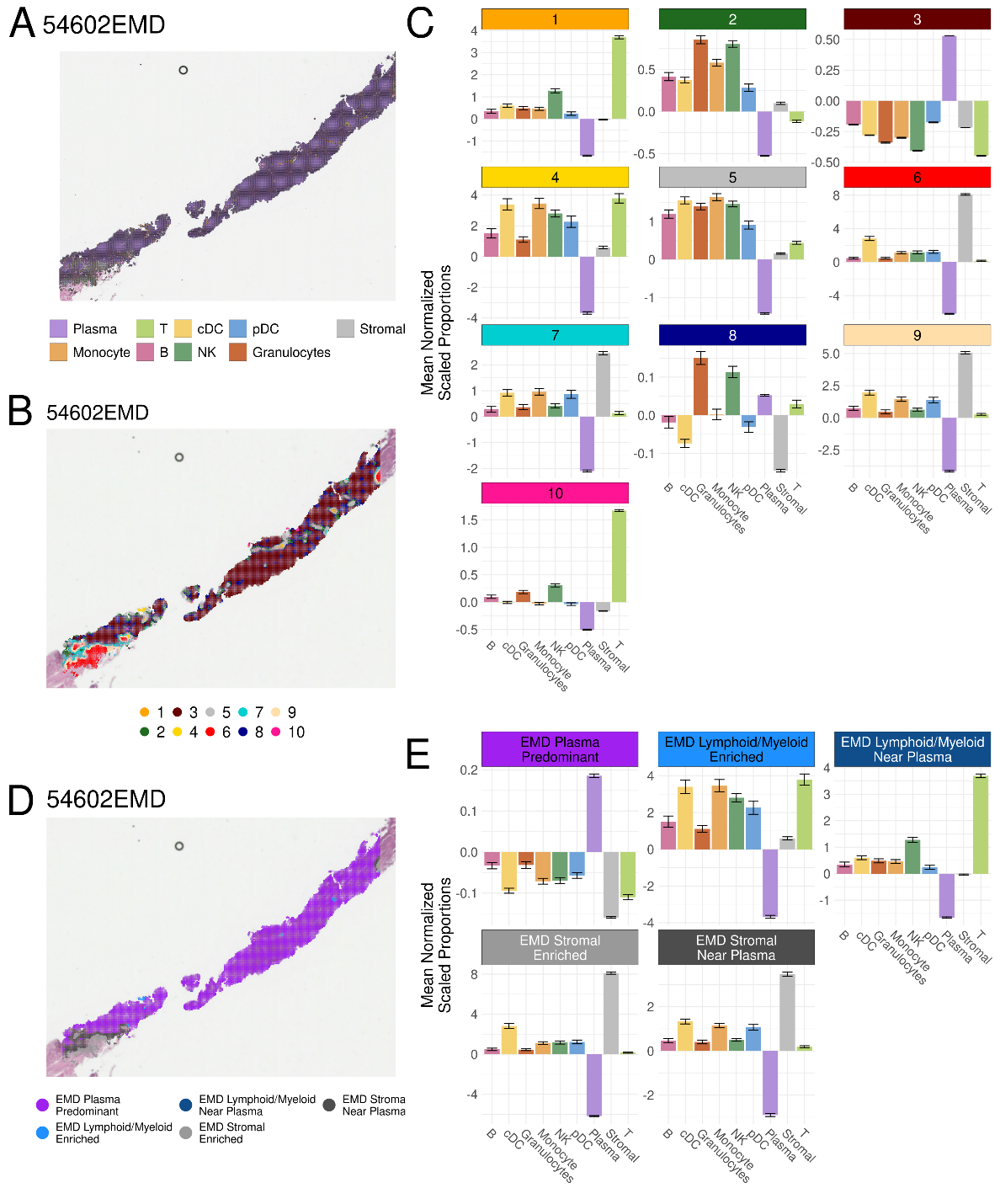

**Supplemental Figure 20. Neighborhood annotation in the extramedullary myeloma Visium HD cohort.** Cells in the extramedullary (EM) Visium-HD cohort underwent the same K-nearest neighbor (KNN) clustering analysis as the bone marrow cohorts. **(A)** Representative spatial dot plot showing the cell type annotations in 54602EMD. **(B)** Representative spatial dot plot displaying the unsupervised KNN clusters. For each voxel, the proportion of cell types within a 50 µm radius was quantified and used for KNN clustering of voxels by proximal cell composition. **(C)** Bar plots depicting the relative abundance of cell types in the twelve KNN clusters. **(D)** Representative spatial dot plot displaying the neighborhood annotations. KNN clusters were annotated based on the enriched cell types as displayed in C. **(E)** Bar plots depicting the relative abundance of cell types in the five annotated neighborhoods.

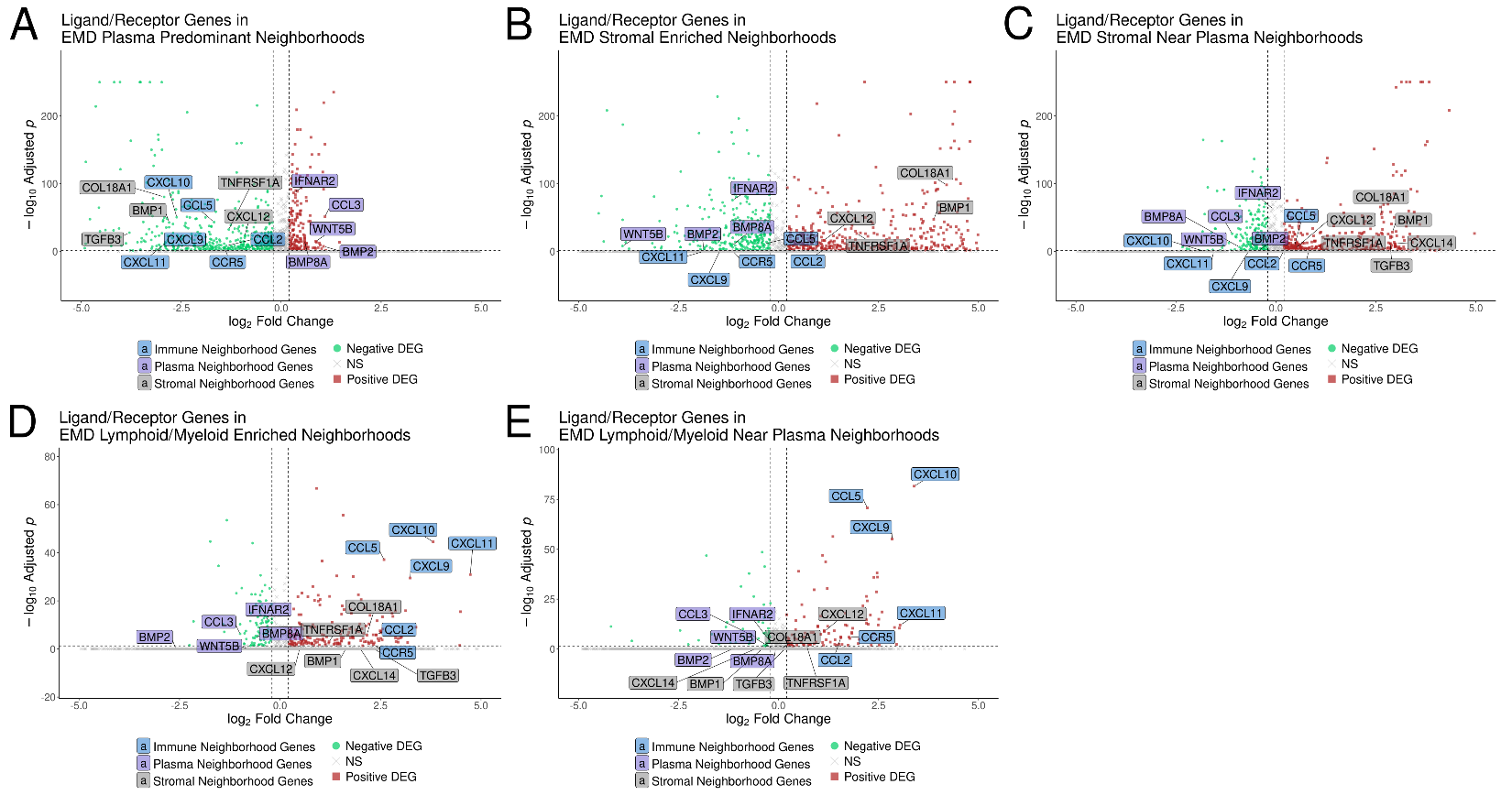

**Supplemental Figure 21. Plasma, immune, and stromal neighborhoods have specific expression of ligand and receptor genes. (A-G)** Volcano plots for each neighborhood in the extramedullary Visium HD cohort testing for enrichment for ligand and receptor genes. Differential expression was tested using a hurdle model as implemented by the MAST method in Seurat’s FindMakers function with a significance cutoff of multiple comparisons adjusted p value < 0.05. Gene labels are colored by whether they were consistently observed to be enriched in the immune neighborhoods, stromal neighborhoods, or in the plasma-predominant neighborhoods.
